# Integrative transcriptomic analysis identifies miR-642a-5p as a regulator of *POFUT1* expression in colon cancer

**DOI:** 10.1101/2025.11.03.686298

**Authors:** Oumaima Mazour, Mouna Ababou, Bouabid Badaoui, Agnès Germot

**Affiliations:** LABCiS (Laboratory of Agroresources, Biomolecules and Chemistry for Health Innovation), UR 22722, University of Limoges, F-87000 Limoges, France; Laboratory of Biodiversity, Ecology and Genome, Department of Biology, Faculty of Sciences, University Mohammed V, Rabat 10100, Morocco; African Sustainable Agriculture Research Institute (ASARI), Mohammed VI Polytechnic University (UM6P), Laâyoune 70000, Morocco

**Keywords:** Biomarker, Colorectal cancer, Gene regulation, Machine learning, miRNA, Multiple linear regression

## Abstract

**Background:** Colorectal cancer (CRC) is one of the most common and deadly cancers worldwide, underscoring the urgent need for novel biomarkers and therapeutic targets. Protein *O*-fucosyltransferase 1 (POFUT1) is increasingly recognized as an oncogenic driver in CRC, but the upstream regulatory mechanisms responsible for its overexpression remain poorly understood.

**Objective:** We aimed to identify a reproducible miRNA signature potentially regulating *POFUT1* in colon cancer by combining multiple computational approaches and intersecting their outcomes to retain the most significant candidates and to validate biologically the most promising miRNA to confirm its regulatory potential.

**Methods:** We implemented an integrative, multi-step bioinformatics framework that combined multiple linear regression with stepwise selection, penalized regression models (LASSO and Elastic Net), and Random Forest analysis for non-linear feature evaluation. miRNAs showing a consistent inverse association with *POFUT1* across these complementary methods were retained as final candidates. The most significant miRNA was then validated in vitro by mimic transfection, real-time quantitative PCR, and dual-luciferase reporter assays performed in colon cancer cell lines.

**Results:** Five miRNAs (miR-92b, miR-484, miR-574, miR-642a, and miR-3940) were consistently identified as negative contributors to *POFUT1* expression across all analytical approaches. Among these, miR-642a exhibited the strongest negative correlation (r = - 0.44) with *POFUT1*. It was therefore selected for biological validation to confirm its regulatory interaction. Overexpression of miR-642a-5p through mimic transfection significantly reduced *POFUT1* transcript levels by approximately 46 % in HCT 116, 50 % in SW480, and 22 % in SW620 cells compared with controls. Dual-luciferase reporter assays confirmed direct binding of miR-642a-5p to two predicted sites within the *POFUT1* 3′UTR, resulting in a decrease of luciferase activity by 40-50 %.

**Conclusion:** This study combines bioinformatics screening based on multiple linear regression and machine learning models, with experimental validation to identify miRNAs regulating *POFUT1* expression in colon cancer. This integrative strategy allows the identification of a miRNA signature related to *POFUT1* and provides novel insights into its post-transcriptional control. It may serve as a foundation for future diagnostic or therapeutic strategies in CRC.

## 1. Introduction

Colorectal cancer (CRC) is one of the most significant global cancer burdens, ranking as the third most diagnosed cancer and the second leading cause of cancer related deaths worldwide. According to the GLOBOCAN 2022 estimates, 1,926,118 new cases of CRC were diagnosed, accounting for 9.6 % of all cancers, with over 903,859 deaths, representing 9.3 % of total cancer deaths for both sexes combined [1]. The global burden of CRC is projected to increase, with an estimated 3.2 million new cases and 1.6 million deaths in 2040, representing a significant rise in the global prevalence [2]. In recent years, increasing incidence among young adults (ages 25-49) has been observed across several high-income countries such as Australia, Canada, UK-England, France, Germany, and USA [3]. The pathogenesis of CRC is a complex and multifactorial process influenced by a combination of genetic alterations, lifestyle, and environmental factors [4]. Advances in precision medicine and high-throughput molecular profiling have significantly improved our understanding of CRC biology, leading to the identification of novel biomarkers and potential therapeutic targets [5].

Among various molecular regulators implicated in cancer, protein *O*-fucosyltransferase 1 (POFUT1) has gained increasing attention in the past decade [6]. The *POFUT1* gene, located on region q11.21 of chromosome 20, encodes a 388-amino acid protein conserved in animals [7–9]. *POFUT1* has been shown overexpressed in oral cancer [10], breast cancer [11,12], hepatocellular carcinoma [13], gastric cancer [14,15], lung cancer [16], colorectal cancer [17,18], oesophageal cancer [19], glioblastoma [20], pancreatic cancer [21], head and neck squamous cell carcinoma [22] and salivary gland carcinoma [23]. POFUT1 is the only enzyme involved in catalyzing the transfer of fucose from GDP-L-fucose to the hydroxyl group of serine (S) or threonine (T) residue within the C^2^XXXX<u>S</u>/<u>T</u>C^3^ consensus sequence, found in epidermal growth factor-like domains (EGF-LDs) of around 100 predicted or characterized glycoproteins [7,24,25]. Among its known targets, Notch receptors have been extensively studied due to the high numbers of EGF-LDs with *O*-fucose modification on their NECD, essential for elongation of the *O*-glycan, necessary to interaction with Notch ligands and the subsequent activation of the Notch canonical pathway [26,27]. In CRC, *POFUT1* is overexpressed in tumor tissues compared to healthy adjacent ones and is associated with tumor sites and pathologic stages [17]. Colon cancer cell lines knockdown for *POFUT1* show decrease cleaved NICD (Notch IntraCellular Domain) levels, and subsequent attenuation of the NOTCH signaling pathway [18]. Their subcutaneous grafts in nude mice significantly reduce tumor size (volume and weight) and induce less liver metastases, compared to control mice. However, despite its oncogenic role, the underlying mechanisms leading to *POFUT1* overexpression in CRC remain poorly understood, which could hinder the development of targeted therapies.

One potential regulatory mechanism that may regulate *POFUT1* could involve microRNAs (miRNAs) [6]. They are small, non-coding RNA, containing approximately 20 to 24 nucleotides. They play a crucial role in post-transcriptional gene regulation by direct binding to complementary sequences on target mRNAs (principally to the 3’UTR), resulting in mRNA degradation or translational inhibition at the protein level [28,29]. miRNAs are involved in early embryonic development and organogenesis. They participate in various biological processes such as cell proliferation, differentiation and apoptosis. Many miRNA gene families are evolutionary conserved among species, which underscores their functional importance. Dysregulation of miRNAs has been linked in humans to several diseases, including cancer, indicating their potential importance in the oncogenic processes [30,31]. In CRC, some miRNAs act as oncogenes by promoting tumor progression, while others function as tumor suppressors, inhibiting cancer growth [32–34]. Their dysregulations offer a broad potential of diagnostic biomarkers and therapeutic targets. Numerous studies have proposed prognostic and predictive CRC markers (for a detailed list, see [35]). Among others, miR-21, expressed at high levels in colon carcinoma cells, is associated with poor survival in CRC patients and a limited response to 5-FU-based adjuvant chemotherapy [36]. The expression levels of let-7 miRNAs, low during developmental processes and more elevated in differentiated tissues, are downregulated in CRC [37]. They are recognized as tumor-suppressive miRNAs and play important roles in regulating the response to anti-epidermal growth factor receptor (EGFR) therapies. In CRC patients with some *KRAS* mutations, a high level of let-7a identifies those who may still benefit from EGFR inhibition therapies [38].

Given the involvement of *POFUT1* in CRC pathogenesis, a deeper understanding of its regulatory mechanisms could unveil novel diagnostic and prognostic biomarkers. Identifying miRNAs that modulate *POFUT1* expression could provide new insights into post-transcriptional gene regulation in CRC. To address this research gap, we propose an integrative bioinformatics framework combining high-throughput transcriptomic data analysis, multiple linear regression (MLR), and machine learning (ML) models to identify miRNAs associated with *POFUT1* expression. A miRNA signature including five strong candidates (miR-92b, miR-484, miR-574, miR-642a, and miR-3940) was identified. Amongst them, miR-642a-5p consistently emerged as the most significant candidate across all analytical frameworks. To verify this computational prediction, we conducted biological validation experiments using normal and carcinoma colon cell lines. The miR-642a-5p was established through its direct binding to *POFUT1* 3′UTR as the first post-transcriptional regulator of *POFUT1* expression, no longer performing its function properly in colon cancer.

## 2. Materials and methods

### 2.1 Bioinformatics analysis

#### 2.1.1 Data acquisition and preprocessing

*POFUT1* and miRNA expression data for colon adenocarcinoma (COAD) were obtained from The Cancer Genome Atlas (TCGA) (https://www.cancer.gov/tcga). Data retrieval and preprocessing were conducted using the TCGAbiolinks package in R (version 4.3.0, https://www.R-project.org/) [39]. For extraction of *POFUT1* expression data, we selected the “Transcriptome Profiling” category with an experimental strategy of “RNA-Seq” and the workflow type “STAR-Counts”. For miRNA expression, data were also collected from the “Transcriptome Profiling” category but using the “miRNA-Seq” experimental strategy and the “miRNA Expression Quantification” for the data type. All datasets were downloaded and processed to remove technical variability. Normalization procedures were applied, followed by a log2 transformation to stabilize the variance across expression levels. Differential expression analysis was performed using the DESeq2 package [40]. To identify highly significant downregulated miRNAs in the COAD cohort, we applied a threshold of log2 Fold Change (FC) < - 2 and adjusted p-value < 0.001, thereby focusing subsequent analyses on the most biologically relevant and significant candidates for a direct explanation of *POFUT1* increase.

#### 2.1.2 Multiple linear regression

To identify a minimal subset of miRNAs providing optimal predictive power, rows containing missing values (NA) were removed before modeling. A MLR model was used with the lm() function in R and the stepwise variable selection strategy based on Akaike Information Criterion (AIC) framework [41] via the stepAIC() function from the MASS package [42]. It was fitted with *POFUT1* expression as the dependent variable and selected miRNAs as independent predictors. Model performance was evaluated through metrics including R^2^, adjusted R^2^, Root Mean Square Error (RMSE), residual standard error (RSE), and F-statistics. To validate assumptions, we examined diagnostic plots: residuals vs fitted values, Q-Q plots, and Cook’s distance.

### 2.1.3 Machine learning

To improve feature selection and capture potential nonlinear interactions, we applied LASSO (Least Absolute Shrinkage and Selection Operator) [43], Elastic Net [44] and Random Forest (RF) regressions as well as the RF classifier [45] implemented in R packages. LASSO regression was applied using a 10-fold cross-validation to determine the optimal penalization parameter (λ). The dataset was split into training (80 %) and testing (20 %) subsets to validate predictive performance. RMSE and R^2^ were calculated on the test set to assess model accuracy. The LASSO regularization technique identified a sparse set of miRNAs with non-zero coefficients, suggesting their importance in predicting *POFUT1*. For robustness, Elastic Net regression was also conducted, enabling a comparison of the miRNA selection consistency under different regularization constraints. RF regression model was constructed, using 500 trees, to account for non-linear interactions between miRNAs and *POFUT1*. Feature importance scores were calculated and used to rank miRNAs based on their contribution to the performance of model prediction. miRNAs were further categorized into high, medium, or low importance groups based on importance score quartiles. The accuracy of the model was evaluated using R^2^ and residual diagnostics.

#### 2.1.4 Calculation of correlation coefficients

To refine feature selection, correlation analysis between selected miRNAs and *POFUT1* expression levels was performed using Pearson correlation coefficients, computed with the cor() function in R. Only significant and negatively correlated miRNAs (r < - 0.1) were retained for further analysis.

#### 2.1.5. Selection of common miRNAs

To identify miRNAs most influencing *POFUT1*, we integrated results from the four comparatives analyses: MLR selected miRNAs, ML selected miRNAs (LASSO/Elastic Net), Random Forest selected miRNAs, and miRNAs negatively correlated with *POFUT1* (r < - 0.1). A Venn diagram was generated to determine the overlap among these selection processes, using library (VennDiagram) in R.

### 2.2 Experimental validation

#### 2.2.1 Cell lines and culture conditions

The normal human colonocyte-derived cell line CCD 841 CoN and the colon cancer cell lines SW480, SW620, and HCT 116 were obtained from the American Type Culture Collection (ATCC, Manassas, VA, USA). CCD 841 CoN cells were maintained in Eagle’s Minimum Essential Medium (EMEM, ATCC) and cancer cell lines were cultured in RPMI-1640 medium (Gibco, Thermo Fisher Scientific, Waltham, MA, USA) with 10 % Fetal Bovine Serum (Gibco, Thermo Fisher Scientific) and 0.5 % antibiotics (100 U/mL penicillin and 100 µg/mL streptomycin) (Gibco, Thermo Fisher Scientific). All cells were maintained at 37 °C in a humidified atmosphere containing 5 % CO2. These cancer cell lines represent distinct stages of colon cancer progression and harbor different mutational profiles.

#### 2.2.2 Transfections with miRNA mimics

Cells were seeded in 6-well plates and transfected 24 h later with 10 nM miRVana™ miR-642a-5p mimic (assayID: MC11477) or negative control (NC) miRNA (reference: 4464058) (Thermo Fisher Scientific). Transfections were performed using INTERFERin® reagent (Polyplus, Ozyme, Saint-Cyr-l’École, France) in Opti-MEM reduced serum medium (Gibco, Thermo Fisher Scientific) according to the manufacturer’s protocol. Cells were incubated for an additional 48 h before RNA extraction. All transfections were conducted in triplicate, in at least three independent experiments.

#### 2.2.3 RNA isolation and real-time qPCR

Total RNA was extracted using the RNeasy Mini Plus Kit (Qiagen, Hilden, Germany) following the manufacturer’s guidelines. RNA quality and quantity were evaluated by using NanoDrop™ spectrophotometer (Thermo Fisher Scientific). A total of 1 µg of RNA and miRNA were reverse transcribed to cDNA using the high-capacity cDNA reverse transcription kit (Applied Biosystems, Thermo Fisher Scientific). Real-time quantitative PCR (RT-qPCR) was performed using TaqMan™ universal PCR master mix and specific TaqMan™ primer assays (Thermo Fisher Scientific). Expression levels of miR-642a-5p (assayID: 001592) were normalized to RNU48 (Small nucleolar RNA, C/D box 48) (assayID: 001006) as an internal control, and *POFUT1* expression (assayID: Hs00382532_m1) was normalized to B2M (β-2-microglobulin) (assayID: Hs00187842_m1). Relative expression levels were calculated using the 2^-ΔΔCt^ method [46]. For miRNA expression, to allow direct comparison with *in silico* transcriptomic data, the resulting fold-change values were log2-transformed. Each sample was analyzed in triplicate, and all experiments were repeated independently at least three times.

#### 2.2.4 Luciferase reporter assay

To evaluate direct binding between miR-642a-5p and *POFUT1* 3′UTR, a dual-luciferase reporter assay (Promega, Madison, WI, USA) was conducted. Two distinct fragments of the *POFUT1* 3′′UTR, each containing one of the two predicted miR-642a-5p binding sites according TargetScan, were selected for analysis. For each region, both wild type (WT) and mutant (MUT) versions were designed. After annealing, each oligonucleotide pair was cloned between the *Sac*I and *Xba*I restriction sites of the pmirGLO Dual-Luciferase miRNA Target Expression Vector (Promega), generating four constructs: pmirGLO-*POFUT1*-WT1, pmirGLO-*POFUT1*-MUT1, pmirGLO-*POFUT1*-WT2, and pmirGLO-*POFUT1*-MUT2. All inserts were confirmed by sequencing. HCT 116 cells were seeded in 96-well plates (5 × 10^4^ cells/well) and allowed to adhere for 24 h prior to transfection. Cells were co-transfected with 100 ng of either WT or MUT reporter construct together with 10 nM miR-642a-5p mimic or NC miRNA using Lipofectamine 3000 (Thermo Fisher Scientific), following the manufacturer’s instructions. Luciferase activity was measured 24 h post-transfection using the Dual-Glo Luciferase Assay System (Promega). Firefly luciferase activity was normalized to *Renilla* luciferase. All transfections were performed in triplicate, in at least three independent experiments.

## 3. Results

### 3.1 Selection of miRNAs associated to *POFUT1* using multiple linear regression

To investigate how miRNAs might regulate *POFUT1* expression in colorectal cancer, we developed a predictive approach using multiple linear regression (MLR) and data from colon adenocarcinoma (COAD) samples available in GDC (Genomic Data Commons) database (https://portal.gdc.cancer.gov/) of the USA National Cancer Institute. Differential expressions of 1410 miRNAs were calculated (Fig. 1a). One hundred of fifteen presented an overexpression > 2 and 217, a downregulation < - 2, with FDR < 0.001. After cleaning the dataset to include complete cases required for MLR, we only considered 67 miRNAs on 238 patient samples. Due to the limited sample size for rectum adenocarcinoma (READ) – despite 83 miRNAs meeting the conditions, only 20 samples were complete – a similar approach could not be carried out. Stepwise variable selection based on AIC was used to optimize performance by including only the most informative predictors. The final model showed strong prediction, explaining approximately 68.1 % of the variance in *POFUT1* expression (R^2^ = 0.681; adjusted R^2^ = 0.6309) (Fig. 1b). The RMSE was 0.66, and the RSE was 0.7394, indicating accurate prediction and relatively low variability in the residuals. A substantial improvement in AIC was observed from - 82.15 in the full model to - 94.07 in the optimized version, suggesting a better model fit with fewer predictors. The F-statistic was highly significant (p < 2.2e-16), confirming that the selected miRNAs collectively contributed meaningfully to the variance in *POFUT1* expression. Statistical diagnostics confirmed that the model satisfied linear regression assumptions. Residuals were randomly and symmetrically distributed around zero in the residual vs predicted plot, supporting linearity and homoscedasticity (Fig. 1c). The Q-Q plot showed residuals closely following a normal distribution, with only slight deviations in the tails (Fig. 1d). Furthermore, Cook’s Distance analysis revealed no highly influential data points, indicating that the model is not disproportionately affected by a small number of data points (Fig. 1e). These evaluations support the high predictive accuracy of the MLR suggesting that 29 miRNAs contribute to regulation of *POFUT1* expression in patients from the COAD cohort.

**Fig. 1.**
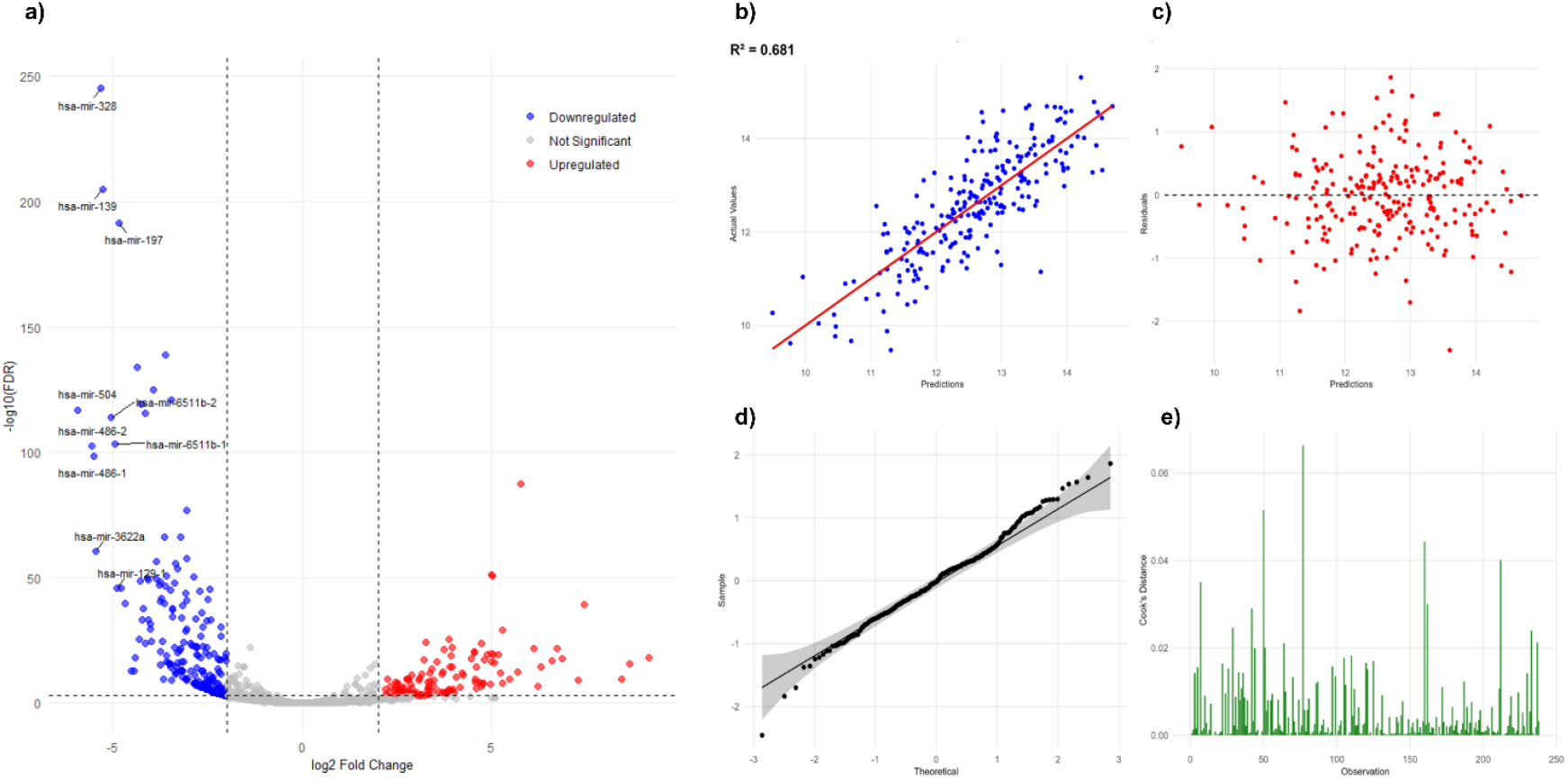
Volcano and diagnostic plots for selection, performance and assumptions of MLR. (**a**) Volcano plot shows fold change (FC) and statistical significance (FDR) of individual miRNA differential expressions between tumor and normal tissues, for patients of the COAD cohort. Blue and red dots represent negatively and positively regulated miRNAs, respectively. Some of the miRNAs, particularly downregulated, were highlighted. (**b**) Actual vs predicted *POFUT1* expression plot shows strong linear correlation between observed and predicted values. (**c**) Residuals vs predicted plot indicates an absence of structural patterns in the residuals and supports homoscedasticity. (**d**) Q-Q plot of residuals confirms normal distribution of errors, with minor deviation at the tails. (**e**) Cook’s Distance plot demonstrates the low influence of individual data on the regression fit

Among them, 13 miRNAs displayed negative regression coefficients, suggesting that increased expressions of these miRNAs were associated with lower levels of *POFUT1* (Table 1). According to their β coefficient, hsa-miR-92a-2 (β = - 2.7696) could be the main miRNA contributor to *POFUT1* expression; however, its p-value (0.1012) was not significant. Among the most significant results, we found hsa-miR-92b (β = - 0.3255; p < 2.59e-06), hsa-miR-3127 (β = - 0.2310; p < 0.0018) and hsa-miR-642a (β = - 0.1457; p < 0.0004).

**Table 1.**
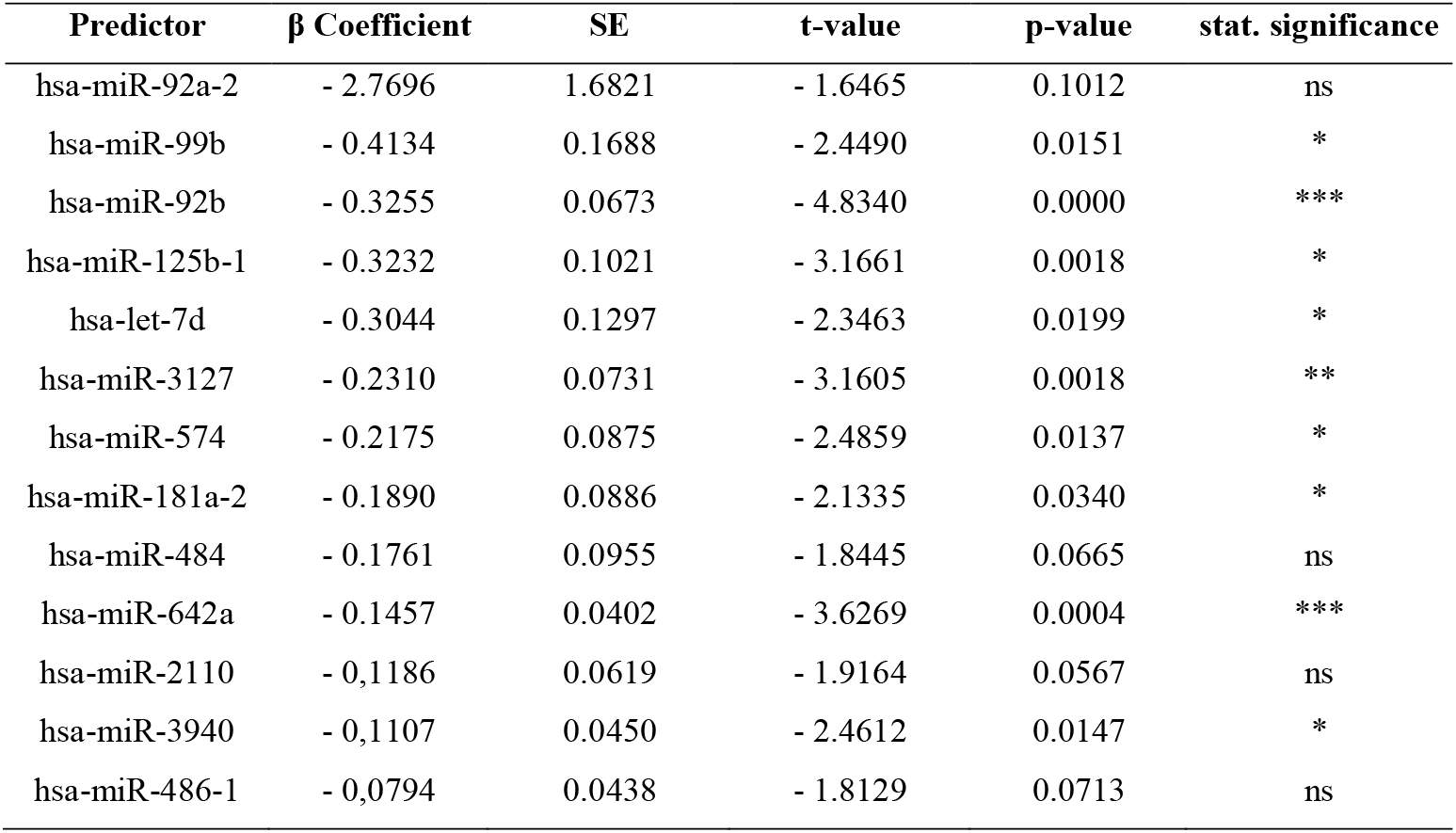
miRNAs with negative coefficients and associated with *POFUT1* in MLR.

| Predictor | $\beta$ Coefficient | SE | t-value | p-value | stat. significance |
| --- | --- | --- | --- | --- | --- |
| hsa-miR-92a-2 | - 2.7696 | 1.6821 | - 1.6465 | 0.1012 | ns |
| hsa-miR-99b | - 0.4134 | 0.1688 | - 2.4490 | 0.0151 | * |
| hsa-miR-92b | - 0.3255 | 0.0673 | - 4.8340 | 0.0000 | *** |
| hsa-miR-125b-1 | - 0.3232 | 0.1021 | - 3.1661 | 0.0018 | * |
| hsa-let-7d | - 0.3044 | 0.1297 | - 2.3463 | 0.0199 | * |
| hsa-miR-3127 | - 0.2310 | 0.0731 | - 3.1605 | 0.0018 | ** |
| hsa-miR-574 | - 0.2175 | 0.0875 | - 2.4859 | 0.0137 | * |
| hsa-miR-181a-2 | - 0.1890 | 0.0886 | - 2.1335 | 0.0340 | * |
| hsa-miR-484 | - 0.1761 | 0.0955 | - 1.8445 | 0.0665 | ns |
| hsa-miR-642a | - 0.1457 | 0.0402 | - 3.6269 | 0.0004 | *** |
| hsa-miR-2110 | - 0.1186 | 0.0619 | - 1.9164 | 0.0567 | ns |
| hsa-miR-3940 | - 0.1107 | 0.0450 | - 2.4612 | 0.0147 | * |
| hsa-miR-486-1 | - 0.0794 | 0.0438 | - 1.8129 | 0.0713 | ns |

### 3.2 Selection of miRNAs associated to *POFUT1* using machine learning analyses

To complement the MLR analysis and to better capture complex potential non-linear relationships, we applied ML (Machine Learning) models to explore the relationship between *POFUT1* expression and the 67 miRNA predictors from the 238 colon adenocarcinoma samples. Two approaches were employed: LASSO regression for feature selection and Random Forest for identifying variable importance in a non-parametric framework. The LASSO algorithm was applied to the processed dataset. It allows selecting the most predictive subset of features, shrinking the others to zero. We used a train-test split as an evaluation method dividing our dataset in two parts, 80 % as a training set (used to train our model) and 20 % as a testing set (used to evaluate our model). The optimal penalization parameter (λ = 0.0607) was selected via a 10-fold cross-validation to minimize prediction errors (Fig. 2a). On the test set, the LASSO regression achieved a R^2^ of 0.63 and a RMSE of 0.74, indicating accurate prediction with a reduced set of variables (Fig. 2b). The residuals vs predicted plot (Fig. 2c) displayed homogeneous residual distribution around zero, while the Q-Q plot (Fig. 2d) showed minor deviation, only at the tails, from the regression line.

**Fig. 2.**
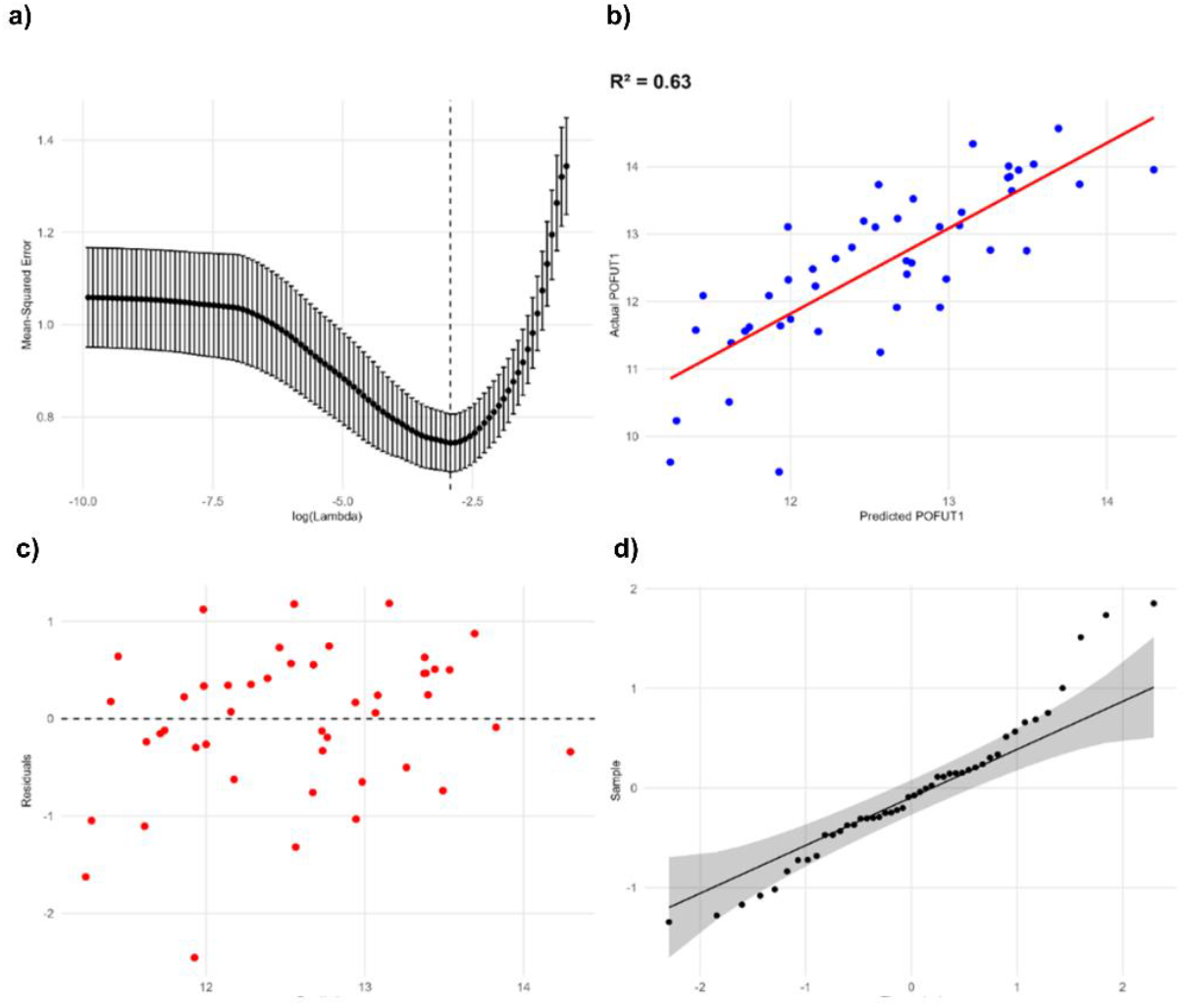
Diagnostic plots evaluating the performance of the LASSO regression method. (**a**) The 10-fold cross-validation curve identifies the optimal penalty parameter (λ). (**b**) Actual vs predicted *POFUT1* expression values show strong linear correlation. (**c**) Residuals vs predicted plot illustrates homoscedasticity and absence of bias. (**d**) Q-Q plot demonstrates normality of residuals, with minor tail deviations

In terms of feature selection, LASSO regression identified a subset of 21 miRNAs most relevant to explain *POFUT1* expression. As a complementary validation, we also applied Elastic Net regression to potentially enlarge the miRNAs dataset identified through LASSO, as this regularization technique minimizes the potential collinearity of the predictive variables. Focusing on negative β coefficients, 9 miRNAs were identified as linked to *POFUT1* overexpression using LASSO, and 4 more, with Elastic Net (Table 2). The top negative contributors were miR-92b (β = - 0.2078 with LASSO / - 0.2135 with Elastic Net), miR-642a (β = - 0.1922 / - 0.1742) and miR-574 (β = - 0.1040 / - 0.1252).

**Table 2.** miRNAs with negative coefficients for LASSO and Elastic Net regularizations, in machine learning MLR.

| miRNA | $\beta$ Coefficient LASSO | $\beta$ Coefficient Elastic Net |
| --- | --- | --- |
| hsa-miR-92b | - 0.2078 | - 0.2135 |
| hsa-miR-642a | - 0.1922 | - 0.1742 |
| hsa-miR-574 | - 0.1040 | - 0.1252 |
| hsa-miR-3940 | - 0.0827 | - 0.0972 |
| hsa-miR-3605 | - 0.0782 | - 0.0788 |
| hsa-miR-484 | - 0.0652 | - 0.0713 |
| hsa-miR-149 | - 0.0328 | - 0.0402 |
| hsa-miR-486-1 | - 0.0290 | - 0.0436 |
| hsa-miR-197 | - 0.0178 | - 0.0649 |
| hsa-miR-3127 | 0 | - 0.0356 |
| hsa-let-7d | 0 | - 0.0067 |
| hsa-miR-370 | 0 | - 0.0033 |
| hsa-miR-375 | 0 | - 0.0022 |

Random Forest is an ensemble of ML methods used for both classification and regression tasks. We trained it with 500 decision trees to capture potential non-linear associations and interactions. The Random Forest algorithm achieved a R^2^ of 0.6, indicating that 60 % of the variance in *POFUT1* expression was explained by the ensemble of miRNA predictors (Fig. 3a). Residual analysis revealed no clear pattern or bias (Fig. 3b). To interpret model output, feature importance scores were calculated, and miRNAs were grouped accordingly into high, medium, and low importance categories based on quartile-pattern boostrapping (Fig. 3c). Among the 23 most influential miRNAs (high category), hsa-miR-92a-1 (25.516), hsa-miR-642a (23.487), hsa-miR-92a-2 (22.055), hsa-miR-92b (15.458), hsa-miR-3940 (12.219) and hsa-miR-3605 (8.168) had the strongest predictive power, suggesting a possible key role in *POFUT1* overexpression. This ranking confirmed the repeated presence of some miRNAs identified in MLR, strengthening the evidence for their potential regulatory roles towards *POFUT1*.

**Fig. 3.**
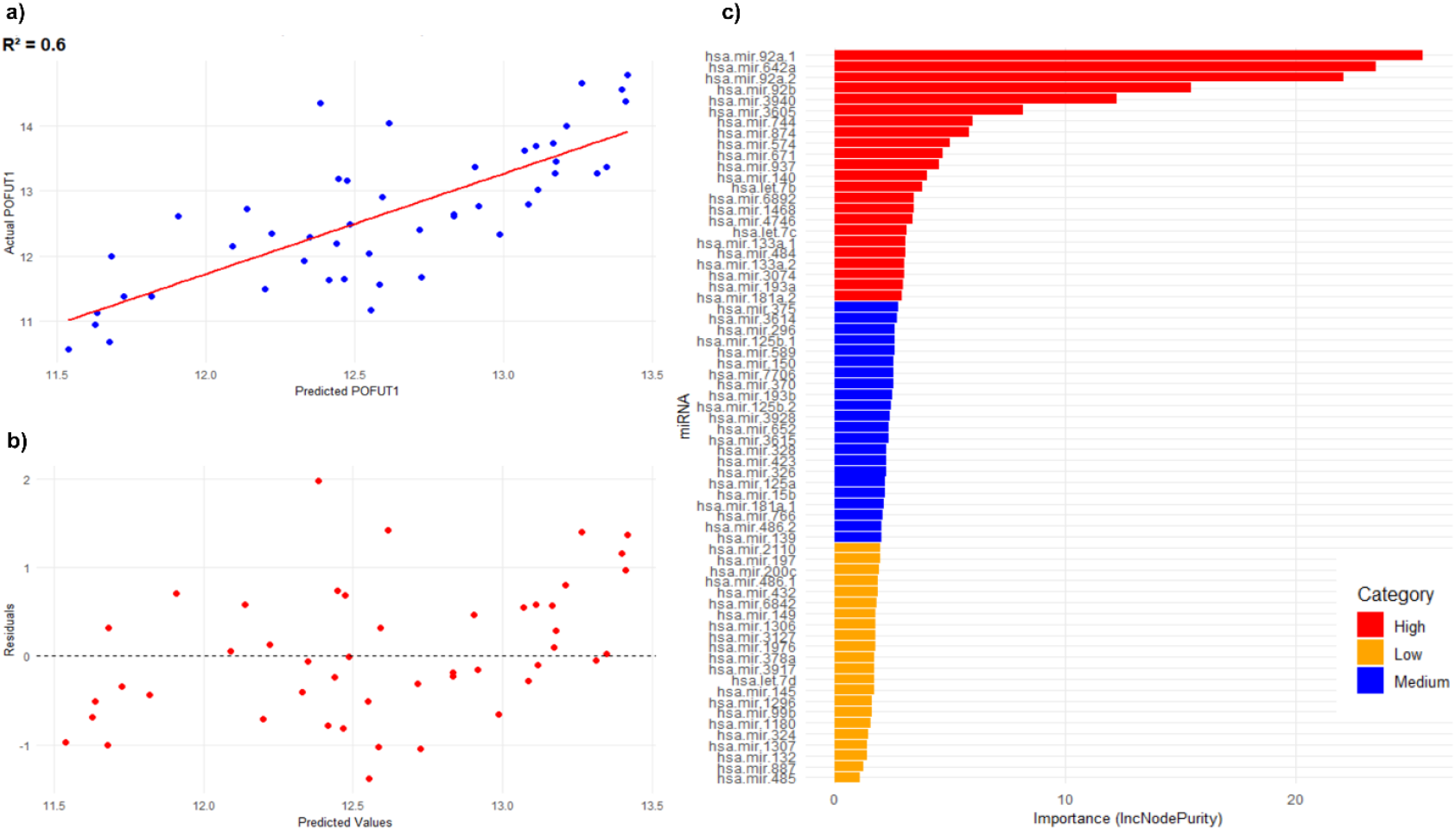
Diagnostic plots evaluating the performance of Random Forest regression and classification of miRNA associated to *POFUT1*. (**a**) Actual vs predicted *POFUT1* expression plot shows strong linear correlation. (**b**) Residuals vs predicted plot illustrates homoscedasticity. (**c**) Bar plots display the 67 miRNAs ordered by decreased scores and categorized in high, medium and low, based on their contributions to node purity

Before investigating correlations with miRNAs, we first confirmed that *POFUT1* remained significantly overexpressed in the subset of patients selected for the analysis, those with complete expression data for both *POFUT1* and miRNAs, after excluding all samples containing missing values (Fig. 4a). The boxplot shows that *POFUT1* is significantly overexpressed in colon tumor tissues compared to adjacent normal ones (p < 0.01).

**Fig. 4.**
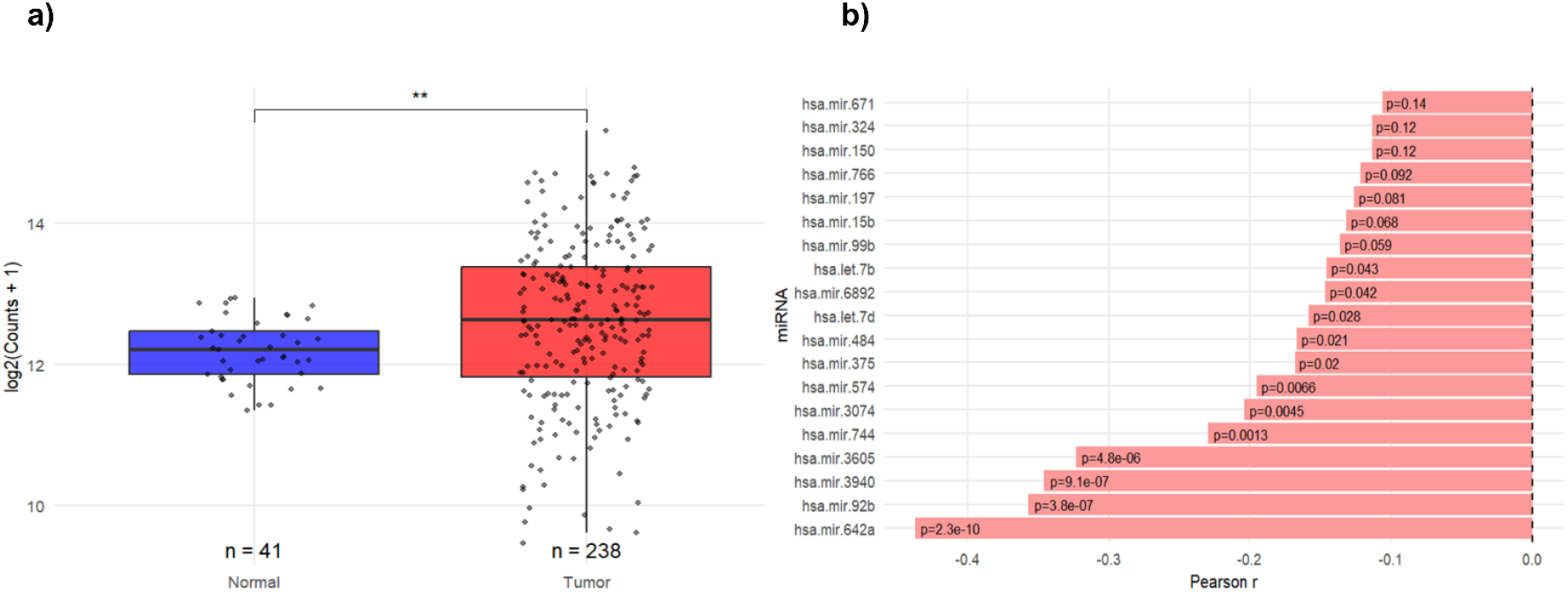
Expression of *POFUT1* in COAD samples and miRNAs classification according to their inverse correlation with *POFUT1*. (**a**) Boxplot comparing *POFUT1* expression levels between tumor samples (n = 238) and adjacent normal tissues (n = 41) from the TCGA-COAD dataset (p < 0.01, Wilcoxon test). (**b**) Bar plots illustrating ordered miRNAs negatively correlated with *POFUT1* expression (r ≤ - 0.1)

To refine candidate selection, we examined the Pearson correlation coefficient (r) for the 67 miRNAs relative to *POFUT1* expression and only retained 19 with r ≤ - 0.1, considering that they may directly target the *POFUT1* transcript and lower its amount in the biological context. It highlighted miR-642a, miR-92b, miR-3940, miR-3605, miR-744, miR-3074 and miR-574. They had the most pronounced inverse relationships with *POFUT1*, supported by statistically significant inverse correlations (p < 0.01) (Fig. 4b). To investigate further the functional relevance of these candidates, we also assessed the correlation between their mature miRNA sequences and *POFUT1* expressions, using the StarBase database (https://rnasysu.com/encori/index.php). Only mature miRNAs that displayed a negative correlation with *POFUT1* in this independent dataset were selected, reinforcing their potential involvement in direct post-transcriptional regulation of *POFUT1*.

### 3.3 Selection of pertinent miRNAs and functional enrichment of their target genes

To extract the most relevant miRNAs, a comparative analysis of miRNA signatures found across MLR, machine learning (LASSO/Elastic Net, Random Forest), and correlation analyses were conducted using a Venn diagram (Fig. 5a) (Supplemental Table 1). The intersection revealed five key miRNAs: hsa-miR-92b, hsa-miR-484, hsa-miR-574, hsa-miR-642a and hsa-miR-3940. Their repeated presence across multiple analytical frameworks argues for their potential role as regulatory miRNAs influencing *POFUT1* expression. To explore KEGG (Kyoto Encyclopedia of Genes and Genomes) pathway enrichment on the union of all predicted targets of these five miRNAs, we submitted these candidates to MIENTURNET (http://userver.bio.uniroma1.it/apps/mienturnet/), which integrates predictions from multiple repositories [47]. From a dataset of 27 to 891 miRNA targets for respectively miR-574-3p and miR-484 (Supplemental Table 2), the analysis revealed five significantly enriched pathways and disease networks (adjusted p-value < 0.05), such as apoptosis, transcriptional dysregulation in cancer, platinum drug resistance, small cell lung cancer, thyroid cancer, and hepatitis C (Fig. 5b). The resulting interaction network (Supplemental Table 3) showed that only hsa-miR-92b-3p and hsa-miR-642a-5p are predicted to target *POFUT1*, as well as other 16 targets (Fig. 5c).

**Fig. 5.**
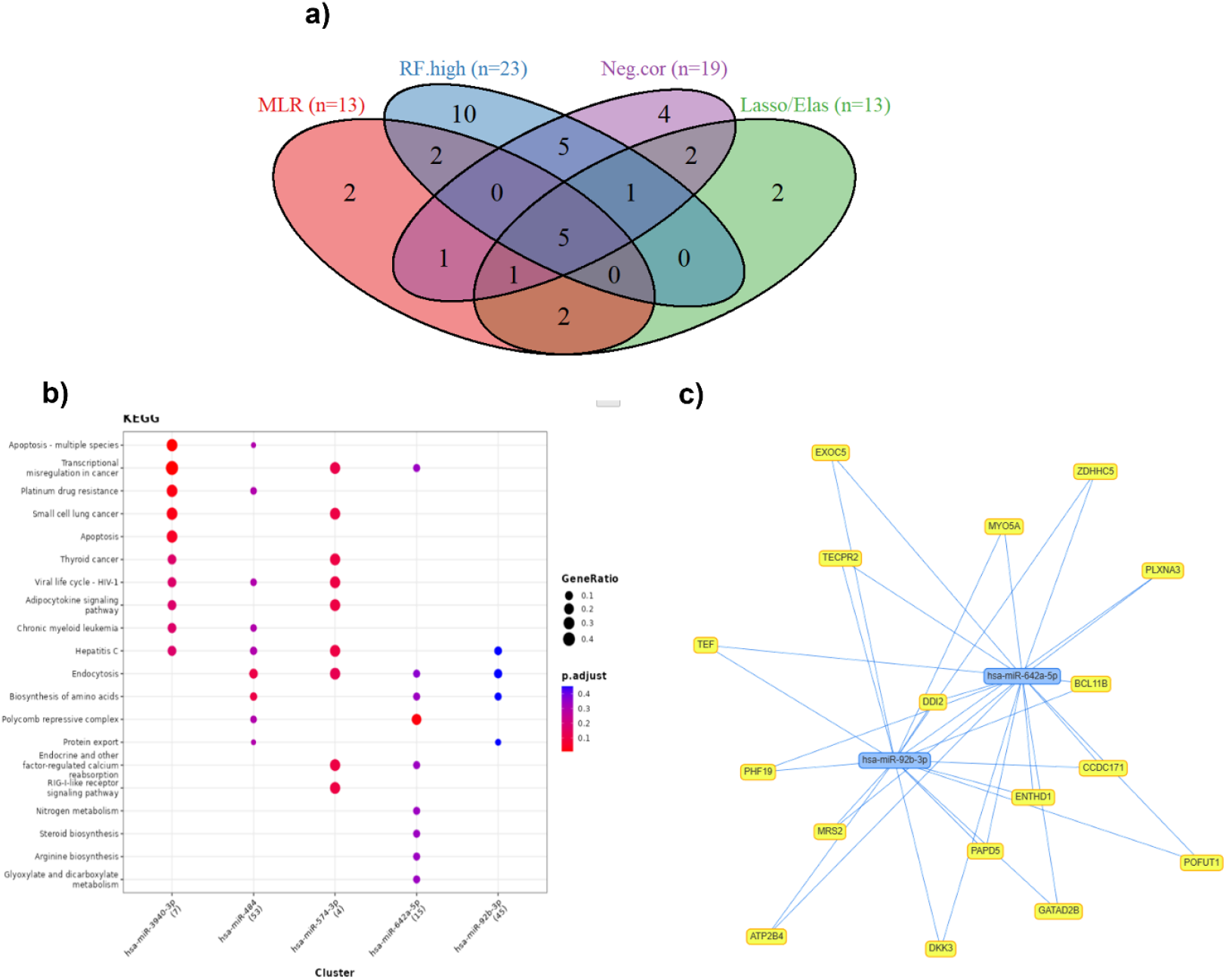
Common miRNA signatures, and molecular interaction, reaction and relation networks. (**a**) Venn diagram of miRNAs obtained from analyses of Multiple Linear Regression (MLR), LASSO/Elastic Net (Lasso/Elas), Random Forest (RF.high), and Pearson correlation coefficient (r ≤ –0.1) with *POFUT1* (Neg.cor). (**b**) KEGG pathway enrichment generated by MIENTURNET from the combined target set of the five common miRNAs. (**c**) miRNA-mRNA interaction network visualized in MIENTURNET, for the two miRNAs targeting *POFUT1*

### 3.4 Expression profiling of *POFUT1* and miR-642a-5p in normal and tumor colon cell lines

Among the five miRNAs signature of *POFUT1* overexpression, miR-642a was selected for biological validation because it exhibited the strongest negative correlation and the most consistent association with *POFUT1* across all computational models. In addition, miR-642a-5p was identified in several prediction and experimentally validated databases as a putative regulator of *POFUT1*. It also showed a pronounced downregulation in the TCGA-COAD dataset compared with normal tissues, reinforcing its potential biological relevance. Endogenous expression levels of *POFUT1* and miR-642a-5p were examined in three colon cancer cell lines (HCT 116, SW480, SW620), representing different stages of tumor progression and metastatic potential. qRT-PCR analysis revealed a marked increase of *POFUT1* mRNA in all tumor cell lines compared with normal colonocyte-derived cell line CCD 841 CoN, as a control (Fig. 6a). *POFUT1* expression increased by 3.6 ± 0.9 in HCT 116, 10.9 ± 1.1 in SW480, and 8.9 ± 2.3 in SW620 relative to control. Expression analysis revealed a consistent and significant downregulation of miR-642a-5p in all colon cancer cell lines relative to CCD 841 CoN used as the reference (Fig. 6b). The mean logFC were - 3.1 ± 0.3 for HCT 116, - 3.2 ± 0.8 for SW480, and - 2.7 ± 0.4 for SW620 (p < 0.01). This decrease displayed an inverse expression pattern relative to *POFUT1* in the same cell lines and closely aligned with the TCGA-COAD data, where miR-642a-5p showed a comparable downregulation with a mean logFC of - 4.3, supporting a negative regulatory relationship between miR-642a-5p and *POFUT1* in colon cancer.

**Fig. 6.**
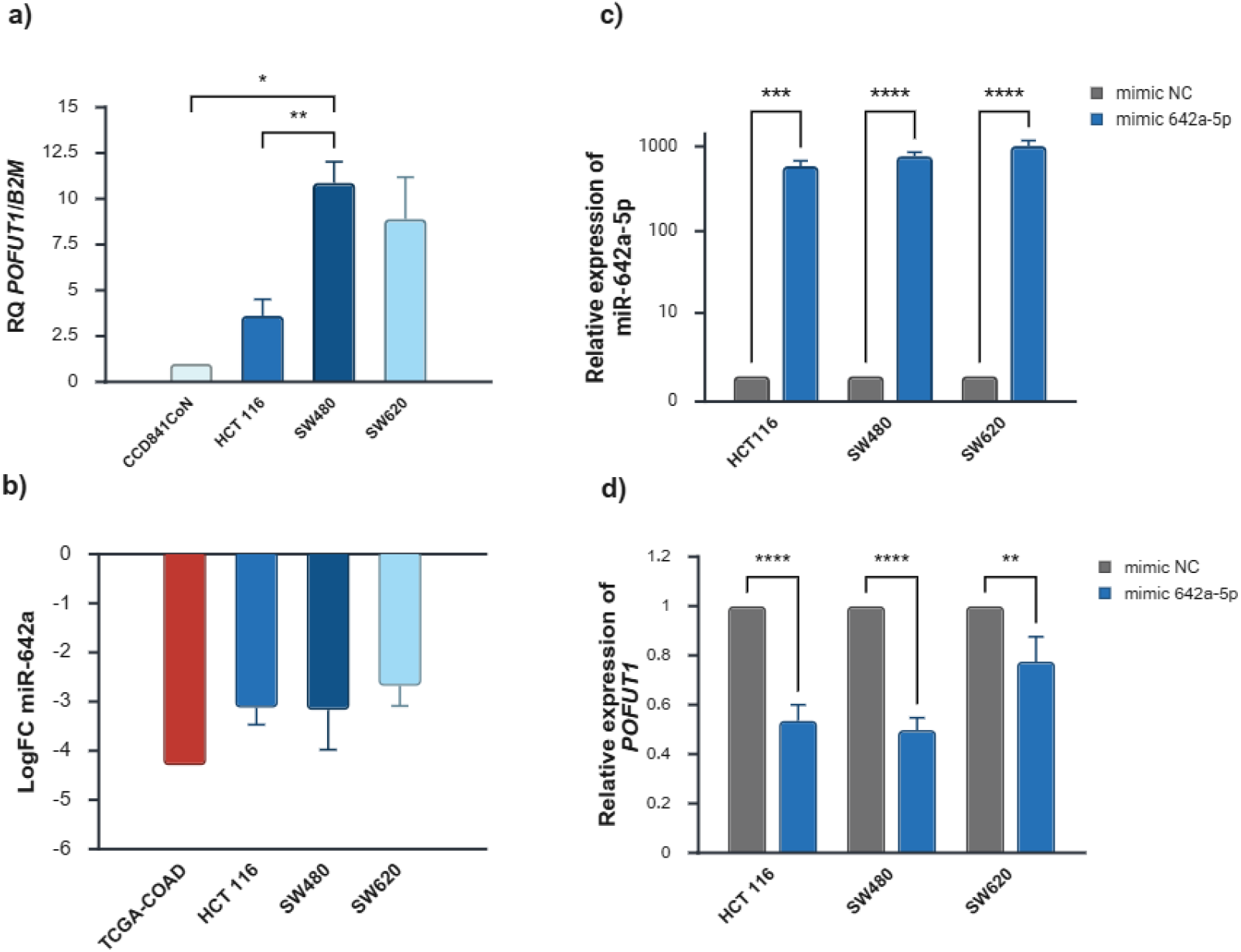
Expressions of *POFUT1* and miR-642a-5p in colon cancer cell lines, before and after transfection by mimic. (**a**) *POFUT1* mRNA expression measured by RT-qPCR in normal (CCD 841 CoN) and cancer colon cell lines (HCT 116, SW480, SW620). Expression values were normalized to *B2M* and expressed as relative quantity (RQ) to CCD 841 CoN. (**b**) Expression of miR-642a-5p in logFC for colon cancer cell lines compared with COAD samples, showing consistent downregulation relative to normal cell/tissue. (**c**) Validation of miR-642a-5p mimic transfection, showing a marked increase in miR-642a-5p expression across all cell lines compared with NC miRNA. (**d**) Effect of miR-642a-5p overexpression on *POFUT1* mRNA levels. Significant reduction of *POFUT1* expression following miR-642a-5p transfection were observed in all tested cell lines. Data are shown as mean ± SD from three biological replicates, except for TCGA-COAD data. Statistical significance was assessed using Welch’s one-way ANOVA. Significance thresholds: p < 0.05 (*), p < 0.01 (**), p < 0.001 (***), p < 0.0001 (****). Created with BioRender

### 3.5 miR-642a-5p mimic reduces *POFUT1* expression

Transfection with the miR-642a-5p mimic resulted in a pronounced elevation of miR-642a-5p expression in all colon cancer cell lines (Fig. 6c). Compared with the negative control (NC), miR-642a-5p levels increased by 587.4 ± 89.5 in HCT 116, 770.2 ± 85.3 fold in SW480, and 1015.7 ± 156.9 fold in SW620, confirming highly efficient mimic transfection and overexpression of this mature miRNA in the cells. Upregulation of miR-642a-5p was accompanied by a significant decrease in *POFUT1* mRNA levels (Fig. 6d). *POFUT1* expression was reduced to 0.54 ± 0.06 (- 46.5 %) in HCT 116, 0.50 ± 0.05 (-50.3 %) in SW480, and 0.78 ± 0.10 in SW620 (-22.5 %), relative to NC set to 1.

### 3.6 Dual-luciferase reporter assay confirms direct binding of miR-642a-5p to *POFUT1* 3′UTR

To confirm the direct interaction between miR-642a-5p and *POFUT1*, two fragments of the *POFUT1* 3′UTR containing the predicted miR-642a-5p binding sites were cloned into the pmirGLO dual-luciferase vector to generate wild type (WT1 and WT2) and corresponding mutant (MUT1 and MUT2) constructs. In HCT 116 cells, co-transfection with the miR-642a-5p mimic markedly reduced the normalized luciferase activity of WT1. The activity significantly decreased from 1.07 ± 0.03 in cells transfected with NC miRNA to 0.49 ± 0.09 in cells transfected with miR-642a-5p mimic (p < 0.001), representing a reduction of approximately 50 % (Fig. 7a). In contrast, the activity of the corresponding mutant construct (MUT1) remained largely unchanged, with values of 1.03 ± 0.06 for NC and 0.91 ± 0.08 for miR-642a-5p mimic, showing that the effect depended on the integrity of the miRNA seed-matching sequence. A similar trend was observed for the second predicted binding site (Fig. 7b). The luciferase signal of the WT2 construct significantly decreased from 1.07 ± 0.02 for NC miRNA to 0.61 ± 0.08 with miR-642a-5p mimic transfection (p < 0.001), while no significant difference was detected for the corresponding mutant (MUT2), which exhibited activities of 1.01 ± 0.03 in NC and 0.93 ± 0.12 with mimic transfection. These results demonstrate that miR-642a-5p directly binds at least to two distinct regions within the *POFUT1* 3′UTR. The loss of regulation role upon mutation of the target sites confirms the specificity of this interaction, providing strong evidence that miR-642a-5p regulates *POFUT1* expression through direct post-transcriptional targeting mechanisms in colon cancer cells.

**Fig. 7.**
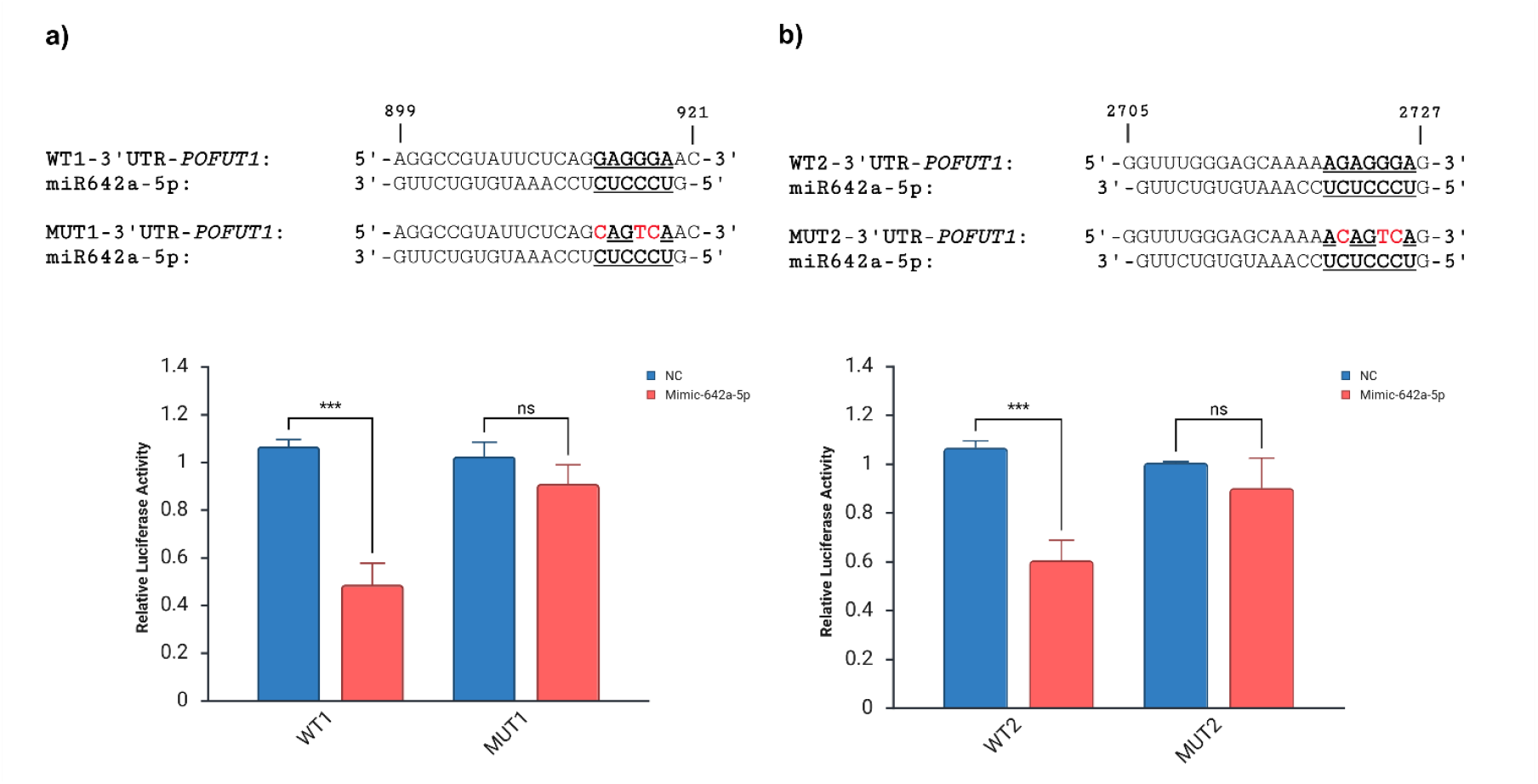
Dual-luciferase reporter assays testing direct binding of miR-642a-5p to two independent regions WT1 (**a**) and WT2 (**b**) of the *POFUT1* 3′UTR. Co-transfection with miR-642a-5p mimic led to a pronounced reduction in luciferase activity for wild type constructs, whereas mutant versions MUT1 (**a**) and MUT2 (**b**) showed no significant difference compared with controls. Numbers above the sequences correspond to positions in the 3’-UTR of *POFUT1* (Genbank accession number, AF375884). Mutations in the targets of miR-642a-5p seed are colored in red. Data are shown as mean ± SD from three biological replicates. Statistical significance was assessed using two-way ANOVA with Tukey’s post hoc test. Significance thresholds: p < 0.001 (***); ns: not significant. Created with BioRender.

## Discussion

POFUT1 has been increasingly recognized as an oncogenic factor in colorectal cancer [6], with different studies demonstrating that it is significantly overexpressed in tumor tissues compared to normal ones [18,48]. *POFUT1* upregulation occurs early during tumorigenesis, being detectable from stage I and persisting into advanced disease stages [17]. Its overexpression promotes tumor progression by enhancing CRC cell proliferation, migration, and invasion, while inhibiting apoptosis. Silencing *POFUT1* through shRNA in colon cancer cells induce cell cycle arrest, decrease migration and invasion capacities, and increase apoptosis, both in vitro and in vivo [18,49]. Therefore, understanding the upstream mechanisms controlling *POFUT1* expression is of considerable interest. In this context, the current study presents a comprehensive exploration of miRNAs that may contribute to post-transcriptional regulation of *POFUT1* in colon cancer, using a multimodal approach that combined classical statistics (multiple linear regression), machine learning (LASSO/Elastic Net regressions and Random Forest), and correlation analysis. Using this analytic framework and in the context of miRNA and mRNA transcriptome data available for the same patients, we consistently identified five miRNAs (miR-92b, miR-484, miR-574, miR-642a, and miR-3940) that were strongly associated with *POFUT1* expression. These miRNAs showed up in all analyses, suggesting that they could play a direct even an indirect role in regulating *POFUT1*. Among these, miR-92b and miR-642a consistently ranked among the most significant predictors of *POFUT1* expression with high statistical significances. They are also listed in the miRNA target prediction databases TargetScan (https://www.targetscan.org/vert_80/) and miRTarBase (https://awi.cuhk.edu.cn/~miRTarBase/miRTarBase_2025/php/index.php) as potential regulators of *POFUT1*.

The miR-642a was retained as a potential negative contributor to *POFUT1* expression in MLR, in machine learning LASSO/Elastic Net regressions, ranked among the top members in the Random Forest and had the strongest negative correlation with *POFUT1* (r = - 0.44). To move beyond prediction and verify whether these computational findings reflected a genuine regulatory mechanism, miR-642a-5p was selected for experimental validation. We showed that an increase of miR-642a-5p expression in colon cancer cell lines led to a significant reduction in *POFUT1* transcript levels, confirming its silencing role at the post-transcriptional level. In fact, *POFUT1* expression decreased by around 46 % in HCT 116, 50 % in SW480, and 22 % in SW620, compared with the negative controls. To further confirm the direct interaction between miR-642a-5p and *POFUT1*, dual-luciferase reporter assays were conducted using two sequence targets of the *POFUT1* 3′UTR. Co-transfection of the miRNA mimic significantly decreased luciferase activity for both constructs, by 40 and 50 % considering the sequences. Therefore, we demonstrated for the first time the existence of a regulatory miRNA for *POFUT1* expression, distinguished after an integrative multi-step bioinformatics approach. Previous studies reported that miR-642a is downregulated in colon cancer as well as in prostate and pancreatic ones (Beveridge et al. 2021; Liu et al. 2022; Shi et al. 2024). Its dysregulation is associated with poor patient outcomes and survival (Ma et al. 2018; Yerukala Sathipati et al. 2022). In colon cancer adenocarcinoma, reduced levels of the miR-642a-5p are linked to increased expression of *MACC1* (Metastasis associated in colon cancer 1), a gene associated with poor prognosis, and to enhanced tumor immune cell infiltration in patients (Liu et al. 2022). The precise mechanism by which this dysregulation would occur is unknown. However, miR-642a, which forms a cluster with miR-642b on chromosome 19, is located in the fragile chromosome region FRA19A. This peculiar location could thus explain a change in the regulation of miR-642a expression in the context of cancer chromosomal rearrangements [50].

Although miR-642a-5p was the only miRNA experimentally validated in this study, the other candidates identified through integrative analysis (miR-92b, miR-484, miR-574, and miR-3940) remain of interest. Their consistent negative association with *POFUT1* expression and their reported tumor-suppressive properties suggest that they could participate in a broader regulatory network influencing *POFUT1* expression and its protein activity. The miR-92b was also significantly downregulated in colon samples of the TCGA-COAD cohort and showed a strong inverse correlation (r = - 0.35) with *POFUT1* expression. It was among the top predictors with significant negative coefficient in MLR, machine learning LASSO/Elastic Net regressions and Random Forest. Exosomal miR-92b level was shown to be significantly downregulated in patients with colorectal carcinomas compared to adenomas, conferring a potential application of this miRNA for an early detection of CRC by liquid biopsy [51]. However, a previous research reported that miR-92b-3p is upregulated in CRC and acts as an oncogene by targeting the tumor suppressor *FBXW7* (F-box and WD40 domain protein 7) and *KLF3* (Kruppel-like factor 3) [52,53]. This contrasting result might be due to differences in tumor subtype and/or sample type used. For example, if in renal cancer, upregulation of miR-92b-3p has been linked to tumor progression [54], in breast cancer, it appears to act as a tumor suppressor by targeting *EZH2* (Enhancer of zeste 2 polycomb repressive complex 2 subunit) [30]. This can be compared to the situation in colon cancer, especially with the selected COAD cohort, where miR-92b-3p may act as a tumor repressor rather than as an oncogenic factor. While miR-484, miR-574, and miR-3940 are not listed in current databases as potential direct regulators of *POFUT1*, they were consistently selected across our analyses as negative contributors of its expression. In particular, miR-484 had a moderate negative coefficient in MLR and a medium feature importance in Random Forest. Its downregulation has been reported in colorectal, gastric, and pancreatic cancers supporting its potential role as a tumor suppressor [55]. Similarly, miR-574 showed a negative association with *POFUT1*. Prior works suggest that miR-574-3p is frequently downregulated in cancer and functions as a tumor suppressor by targeting oncogenic pathways, thereby inhibiting proliferation and metastasis [56–58]. In CRC, reduced expression of miR-574-3p has been linked to enhanced tumor proliferation, metastasis, and therapy resistance, partly through the regulation of *IGF1R* (Insulin-like growth factor 1 receptor) [59]. It also shows significant correlation with tumor size, clinical stage, and lymph node metastasis. Furthermore, miR-574-3p demonstrates diagnostic value, with lower circulating levels effectively distinguishing CRC patients from healthy controls [60]. In the present work, miR-3940 had a strong and significant negative MLR coefficient and was consistently retained as a predictive feature in both LASSO/Elastic Net and Random Forest analysis methods. The mature miR-3940-3p has not been studied in CRC yet; however, in other cancer types, it has been reported to play regulatory roles. In contrast, the mature miR-3940-5p was reported as downregulated in CRC patient serum and tumor tissues, correlating with poor prognosis [61].

Although it is not possible to define a main signaling pathway that would be affected by dysregulation of *POFUT1*-associated miRNAs, it is noteworthy that some of the predicted targets of miR-92b-3p and miR-642a-5p are, like *POFUT1*, overexpressed in CRC. Indeed, *EXOC5* (Exocyst Complex Component 5) is overexpressed in cancerous tissues compared to adjacent healthy ones, promoting tumor growth, partly via the PI3K/AKT (Phosphoinositide 3-kinase/AKT serine-threonine protein kinase) and MAPK (Mitogen-activated protein kinase) signaling pathways [62]. *DDI2* (DNA Damage Inducible 1 Homolog 2), which manages replication stress and maintains genome integrity, is upregulated in the early stages of CRC (stage I–II) and its knockdown reduces proliferative, migratory and invasive capabilities in vitro and in vivo [63]. It acts via the MAPK signaling pathway. Finally, overexpression of *PHF19* (PHD Finger Protein 19) is also linked to tumor progression, especially in epithelial-mesenchymal transition and metastasis [64]. It is associated with poor overall survival in the monitored CRC cohort.

The combination of multiple linear regression and machine learning approaches allowed us to identify miRNAs that explain a significant part of the variation in *POFUT1* expression for colon cancer. These observations raise an important possibility that the lowered levels of those miRNAs in colon cancer may contribute to *POFUT1* overexpression, thereby fueling tumor growth and invasion. Restoring these miRNAs could suppress *POFUT1* overexpression, causing decreased proliferation, migration, and survival of tumor cells. Thus, therapeutic strategies aimed at restoring miR-642a-5p levels could represent a novel approach to target *POFUT1*. In addition, the fact that *POFUT1* overexpression is an early event in CRC [17,18] suggests that dysregulation of these miRNAs may occur as well, early in tumor development. However, it will be important to apply this combinatory approach in larger and more diverse patient cohorts to rule out a possibility of selection bias in the TCGA-COAD cohort. Moreover, as *POFUT1* overexpression is also known to occur in rectum cancer [17], a study of a cohort specific to this tumor location would enable us to determine whether there is a shared signature with the miRNAs identified for the colon and/or whether there are specific rectum biomarkers.

This study provides new insights into the post-translational regulation of *POFUT1* in colon cancer by combining computational prediction and experimental validation. Through an integrative bioinformatics approach, we identified a set of miRNAs inversely associated with *POFUT1* expression and highlighted miR-642a-5p as the most promising candidate. The experimental assays confirmed that this miRNA directly targets two regions of the *POFUT1* 3′UTR, leading to a clear reduction in its expression levels.

These findings establish a solid link between miR-642a-5p and *POFUT1*, suggesting that their combined dysregulation contributes to colon tumorigenesis. Beyond the mechanistic insight, this regulatory axis may hold clinical relevance highlighting miR-642a-5p as both a biomarker of the disease and a potential therapeutic target. The consistency between computational and biological results reinforces our approach. In silico identification of additional miRNAs, miR-92b, miR-484, miR-574, and miR-3940, as potential negative regulators needs, of course, biological confirmations. However, they already support the existence of a coordinated miRNA network modulating *POFUT1* expression and may constitute a distinctive molecular signature of colon cancer, providing a basis for future experimental and translational investigations.

## Supplementary information

Supplementary materials are available.

## Acknowledgements

We would like to thank Master’s students for their contributions to this work as part of their internship, in particular Ms Fatima-Ezzahra Bouktaib, and Dr François Gallet and Pr Tony Lefebvre, for sharing some cell lines. We also acknowledge the TCGA Research Network for providing open access to the COAD dataset, and the developers of the TCGAbiolinks R package for enabling efficient data access from the GDC portal. Finally, we are grateful to the R community for the open-source tools that supported our analyses and visualizations.

## Author contributions

**Oumaima Mazour:** Conceptualization, Methodology, Formal analysis, Investigation, Writing - Original Draft, Visualization. **Mouna Ababou:** Writing - Review & Editing. **Bouabid Badaoui:** Conceptualization, Methodology, Formal analysis, Writing - Review & Editing, Supervision. **Agnès Germot:** Conceptualization, Writing - Review & Editing, Supervision, Project administration, Funding acquisition.

## Data Availability

The following information was supplied regarding data availability:

The dataset was gathered from the TCGA-COAD project via the Genomic Data Commons (GDC) data portal (https://portal.gdc.cancer.gov/), using the TCGAbiolinks R package.

Source code is available upon request. Please contact Oumaima Mazour.

## Ethical approval

Not applicable.

