## Supplemental Table 1 for "Integrative transcriptomic analysis identifies miR-642a-5p as a regulator of *POFUT1* expression in colon cancer"

^*^ co‑last authors

Corresponding author:

Agnès Germot**^*1^**

**Supplemental Table 1** miRNAs selected by multiple feature selection methods used for intersection analysis (Venn diagram)

| **MLR** | **Lasso/Elas** | **RF.high** | **Neg.cor** |
| --- | --- | --- | --- |
| hsa-let-7d | hsa-let-7d | hsa-let-7b | hsa-let-7b |
| hsa-miR-92a-2 | hsa-miR-92b | hsa-let-7c | hsa-let-7d |
| hsa-miR-92b | hsa-miR-197 | hsa-miR-92a-1 | hsa-miR-15b |
| hsa-miR-99b | hsa-miR-149 | hsa-miR-92a-2 | hsa-miR-92b |
| hsa-miR-125b-1 | hsa-miR-370 | hsa-miR-92b | hsa-miR-99b |
| hsa-miR-181a-2 | hsa-miR-375 | hsa-miR-133a-1 | hsa-miR-150 |
| hsa-miR-2110 | hsa-miR-484 | hsa-miR-133a-2 | hsa-miR-197 |
| hsa-miR-3127 | hsa-miR-486-1 | hsa-miR-140 | hsa-miR-324 |
| hsa-miR-3940 | hsa-miR-574 | hsa-miR-181a-2 | hsa-miR-375 |
| hsa-miR-484 | hsa-miR-642a | hsa-miR-193a | hsa-miR-484 |
| hsa-miR-486-1 | hsa-miR-3127 | hsa-miR-484 | hsa-miR-574 |
| hsa-miR-574 | hsa-miR-3605 | hsa-miR-574 | hsa-miR-642a |
| hsa-miR-642a | hsa-miR-3940 | hsa-miR-642a | hsa-miR-671 |
|  |  | hsa-miR-671 | hsa-miR-744 |
|  |  | hsa-miR-744 | hsa-miR-766 |
|  |  | hsa-miR-874 | hsa-miR-3074 |
|  |  | hsa-miR-937 | hsa-miR-3605 |
|  |  | hsa-miR-1468 | hsa-miR-3940 |
|  |  | hsa-miR-3074 | hsa-miR-6892 |
|  |  | hsa-miR-3605 |  |
|  |  | hsa-miR-3940 |  |
|  |  | hsa-miR-4746 |  |
|  |  | hsa-miR-6892 |  |
