## Supplemental Table 2 for "Integrative transcriptomic analysis identifies miR-642a-5p as a regulator of *POFUT1* expression in colon cancer"

^*^ co‑last authors

Corresponding author:

Agnès Germot**^*1^**

**Supplemental Table 2** List of target genes extracted from MIENTURNET for the 5 selected miRNAs

| miRNA-target miRTarBase | | | | |
| --- | --- | --- | --- | --- |
| **hsa-miR-484** | **hsa-miR-92b-3p** | **hsa-miR-642a-5p** | **hsa-miR-3940-3p** | **hsa-miR-574-3p** |
| AARS | AAED1 | ABL2 | AKR1B10 | AMOTL1 |
| AARS2 | ABCA3 | ACADL | ASB6 | C2orf72 |
| ABCD4 | ABCF2 | ADAM33 | ATG9A | CKS2 |
| ABCF1 | ABR | AK3 | BCL2L1 | CLTC |
| ABHD12 | ACAA1 | AKAP2 | BIRC5 | CPLX2 |
| ABHD2 | ACTC1 | ANTXR1 | BLOC1S3 | CUL2 |
| ABL1 | ACTN4 | ANTXR2 | C12orf49 | EGFR |
| ACADS | ADAM10 | ARL11 | C21orf2 | EP300 |
| ACLY | ADAT1 | ARRB1 | CA5B | IGLON5 |
| ACOT9 | AEN | ASIC4 | CCBE1 | IRF4 |
| ACP4 | AGBL5 | ATP2B1 | CYTH1 | KCNK10 |
| ACTA2 | AGMAT | ATP2B4 | EHD4 | LHFPL3 |
| ACTB | AKAP10 | ATP6AP1L | EIF1AD | LHFPL4 |
| ADA | AKAP8 | ATP6V0E1 | GPAT4 | MKRN1 |
| ADAM10 | ALDOA | ATP9A | GPR75 | MUM1 |
| ADAR | ALG14 | BCL11B | HASPIN | PDZD2 |
| ADCK2 | AMD1 | C1orf210 | HRH2 | PEAK1 |
| ADGRE5 | AMIGO1 | C2orf48 | KDM6B | PHEX |
| ADRA1A | ANKIB1 | C6orf223 | KXD1 | PIN1 |
| ADRM1 | ANKRD52 | CALHM5 | LAD1 | PRMT1 |
| AEBP2 | ANP32E | CBX6 | LTBP2 | RAC1 |
| AGK | AP1B1 | CCDC171 | MELTF | RXRA |
| AGL | AP2A2 | CCDC93 | MLLT1 | SMAD4 |
| AGO1 | AP3S2 | CDC42EP4 | MON1B | TGFB1 |
| AGO4 | AP5Z1 | CDH8 | NAV1 | TLNRD1 |
| AHDC1 | APOBEC3F | CELF2 | NPAS1 | TMEM178B |
| AIFM1 | APOLD1 | CEP70 | NRGN | USP31 |
| AKAP13 | APPL1 | CHRM2 | NRXN3 |  |
| AKAP7 | ARF1 | CLDN1 | ORAI2 |  |
| AKT1S1 | ARFGEF2 | CLIC5 | POLA2 |  |
| ALAS1 | ARGFX | CLUAP1 | POU2F1 |  |
| ALKBH5 | ARID1B | CMBL | PPP1R11 |  |
| AMER1 | ARL6IP4 | COL5A3 | PURB |  |
| AMMECR1L | ARNTL2 | CRISPLD2 | QPCTL |  |
| AMOTL1 | ASF1B | CRY2 | RAB11B |  |
| AMPD2 | ASGR2 | CTSV | RILPL1 |  |
| ANAPC1 | ATF7IP | DCLK3 | RPL18A |  |
| ANAPC7 | ATOX1 | DDI2 | RXRA |  |
| ANKRD36 | ATP13A1 | DHCR24 | SCUBE3 |  |
| ANKRD54 | ATP2A2 | DKK3 | SGPP2 |  |
| ANXA11 | ATP2B4 | DNAJC10 | SH2B3 |  |
| ANXA6 | ATP5G3 | DOHH | SIGLEC9 |  |
| AP1B1 | ATP7A | DOK6 | STK10 |  |
| AP1G1 | ATXN1 | DPY19L1 | TERF2 |  |
| AP2M1 | AURKA | DSN1 | TIMELESS |  |
| AP3D1 | B4GALT7 | DUSP4 | TMEM250 |  |
| APEX1 | BAG6 | EEF2 | TMEM41B |  |
| APLP1 | BAK1 | EGLN3 | TRAF6 |  |
| APP | BAZ2B | EIF2B5 | TRPM6 |  |
| APRT | BBX | EMC10 | ZBTB46 |  |
| ARAP1 | BCAT1 | EMX2 | ZNF286A |  |
| ARFGEF2 | BCAT2 | ENDOD1 | ZNF286B |  |
| ARHGDIA | BCKDK | ENTHD1 | ZNF585B |  |
| ARID1A | BCL11B | ERCC1 |  |  |
| ARID3A | BCL2L11 | ETV3 |  |  |
| ARL6IP4 | BICD2 | EXOC5 |  |  |
| ARMC6 | BMP8A | EXTL3 |  |  |
| ASCC2 | BMPR1A | FAAP24 |  |  |
| ASNA1 | BMPR2 | FADS1 |  |  |
| ASXL1 | BPTF | FAM110A |  |  |
| ATF7IP | BRD8 | FAM204A |  |  |
| ATL2 | BRMS1L | FAM20B |  |  |
| ATN1 | BTG2 | FAM210B |  |  |
| ATP11A | C11orf24 | FAM213A |  |  |
| ATP1A1 | C15orf38-AP3S2 | FAM84A |  |  |
| ATP1A2 | C17orf75 | FBRS |  |  |
| ATP2A2 | C1GALT1C1 | FBXO47 |  |  |
| ATP5J | C1orf35 | FCRL4 |  |  |
| ATP6V1F | C21orf91 | FER |  |  |
| ATXN7L3 | C2orf69 | FOXL1 |  |  |
| ATXN7L3B | C5orf24 | FRMD6 |  |  |
| AUNIP | C6orf62 | FRMPD4 |  |  |
| AUP1 | C9orf64 | GALNT10 |  |  |
| AXIN1 | CAD | GATAD2B |  |  |
| B4GALT1 | CAMSAP1 | GJB7 |  |  |
| BACE2 | CAPN15 | GLIPR1L2 |  |  |
| BAG5 | CAPRIN1 | GLUL |  |  |
| BAG6 | CAPZB | GNAZ |  |  |
| BAMBI | CARD6 | GPC5 |  |  |
| BAZ2A | CASD1 | GPR37L1 |  |  |
| BBC3 | CASKIN1 | GPR75 |  |  |
| BBX | CCDC113 | GRID1 |  |  |
| BCAS4 | CCDC171 | GTDC1 |  |  |
| BCAT2 | CCDC186 | GTF2H5 |  |  |
| BCL2L1 | CCDC22 | GTF3C4 |  |  |
| BEND3 | CCNB1 | HEYL |  |  |
| BIRC5 | CCNQ | HLCS |  |  |
| BLMH | CCSER2 | IL2RA |  |  |
| BLOC1S6 | CD180 | IQSEC1 |  |  |
| BMS1 | CD226 | JPH3 |  |  |
| BRAT1 | CD2AP | KCNA7 |  |  |
| BRCA1 | CD69 | KCNK12 |  |  |
| BRD2 | CDC27 | KLHL25 |  |  |
| BRD4 | CDC5L | KLHL9 |  |  |
| BRI3 | CDC6 | KRAS |  |  |
| BSN | CDK16 | KSR2 |  |  |
| BTG1 | CDK5R1 | LARP1 |  |  |
| BUD23 | CDKN1C | LENG9 |  |  |
| C12orf10 | CEP152 | LGMN |  |  |
| C12orf49 | CHST1 | LHX5 |  |  |
| C14orf166 | CIC | LINC00598 |  |  |
| C19orf12 | CIDEC | LYRM2 |  |  |
| C1orf21 | CISD1 | MACC1 |  |  |
| C1QBP | CIT | MAP3K9 |  |  |
| C2orf68 | CLN8 | MAPK14 |  |  |
| C5orf51 | CLTA | MCTP1 |  |  |
| C6orf106 | CNEP1R1 | MDN1 |  |  |
| C8orf44-SGK3 | CNIH1 | MED21 |  |  |
| CABLES1 | CNNM4 | MED4 |  |  |
| CABP4 | CNOT2 | METTL1 |  |  |
| CADM1 | CNOT4 | MEX3A |  |  |
| CAMK2D | COG3 | MRE11 |  |  |
| CAMLG | COPA | MRPS27 |  |  |
| CAP1 | COQ8B | MRS2 |  |  |
| CAPN15 | COX1 | MTA3 |  |  |
| CAPRIN1 | COX20 | MTG1 |  |  |
| CAPZB | CPEB2 | MTRNR2L1 |  |  |
| CASC3 | CPEB3 | MYADM |  |  |
| CASP7 | CPEB4 | MYO5A |  |  |
| CBR1 | CPTP | MYOCD |  |  |
| CBX5 | CREB3L2 | NEDD9 |  |  |
| CBX6 | CRIM1 | NEXMIF |  |  |
| CCDC136 | CSDE1 | NODAL |  |  |
| CCDC47 | CTC1 | NUPL2 |  |  |
| CCDC94 | CTDSPL | OCIAD2 |  |  |
| CCDC97 | CYP20A1 | OLR1 |  |  |
| CCNF | CYP2C19 | ONECUT3 |  |  |
| CCNK | CYTH2 | PAFAH1B2 |  |  |
| CCNT2 | DAB2IP | PALM2 |  |  |
| CCS | DAND5 | PALM2-AKAP2 |  |  |
| CD2AP | DBT | PANK1 |  |  |
| CD2BP2 | DDI2 | PAPD5 |  |  |
| CD3E | DDIT4 | PDCL |  |  |
| CDC25A | DDX3X | PEX5L |  |  |
| CDC34 | DENND2C | PFKM |  |  |
| CDC37L1 | DENND4A | PGM2L1 |  |  |
| CDC42 | DENND4B | PHF19 |  |  |
| CDCA7L | DEXI | PHF7 |  |  |
| CDS2 | DKK3 | PIGR |  |  |
| CDX2 | DLST | PINX1 |  |  |
| CECR2 | DNAAF5 | PITPNM3 |  |  |
| CEP250 | DNAJB12 | PKDREJ |  |  |
| CEP89 | DNAJB9 | PKHD1L1 |  |  |
| CERS5 | DNAJC27 | PLA2G2C |  |  |
| CFL1 | DNAJC30 | PLCXD3 |  |  |
| CHAF1B | DOCK11 | PLEKHG4B |  |  |
| CHCHD2 | DOCK9 | PLXNA3 |  |  |
| CHD4 | DSTYK | POFUT1 |  |  |
| CHMP7 | DUS2 | POU5F1 |  |  |
| CHSY3 | DUSP10 | PPP1R3G |  |  |
| CHTF18 | DUSP5 | PPP6R1 |  |  |
| CKAP5 | DYNC1LI2 | PRKAB1 |  |  |
| CLDN7 | DYNLT3 | PRKAR2A |  |  |
| CLEC16A | E2F3 | PSG4 |  |  |
| CLN8 | EARS2 | PSMB2 |  |  |
| CLPP | EDEM1 | PTCD3 |  |  |
| CLTC | EDF1 | PTGIS |  |  |
| CLUAP1 | EDRF1 | PTPN14 |  |  |
| CLUH | EFNB1 | PTPRG |  |  |
| CMPK1 | EID2B | PXDC1 |  |  |
| CNFN | EIF1 | PYCARD |  |  |
| CNGA4 | EIF2AK1 | QPRT |  |  |
| CNIH4 | EIF2B2 | RAB13 |  |  |
| COG1 | EIF3A | RAB8A |  |  |
| COL18A1 | EIF4EBP1 | RBX1 |  |  |
| COL23A1 | EIF4EBP2 | RGS5 |  |  |
| COPS3 | EIF5A2 | RGS6 |  |  |
| COQ8A | ELOA | RHOB |  |  |
| COX3 | ENTHD1 | RUNX1 |  |  |
| COX5A | EPM2AIP1 | RUSC2 |  |  |
| COX8A | ERGIC2 | SAMHD1 |  |  |
| CPSF7 | ERICH1 | SCUBE3 |  |  |
| CREBRF | ESRP1 | SEMA3E |  |  |
| CRTAP | EVI5 | SETD1A |  |  |
| CTPS1 | EXOC5 | SHISA9 |  |  |
| CTSD | EZH2 | SIX3 |  |  |
| CUL5 | F11R | SKA2 |  |  |
| CUTA | FAM126B | SLC22A6 |  |  |
| CWC22 | FAM129A | SLC25A17 |  |  |
| CYB5R3 | FAM135A | SLC30A3 |  |  |
| CYFIP1 | FAM3C | SLC35B1 |  |  |
| CYSTM1 | FAM46A | SLC39A1 |  |  |
| DCAF1 | FAM49A | SLC5A12 |  |  |
| DCAF7 | FAM91A1 | SMAD2 |  |  |
| DCBLD1 | FAR1 | SMYD1 |  |  |
| DCLRE1B | FASLG | SP9 |  |  |
| DCTN3 | FASN | SPNS1 |  |  |
| DCTN4 | FBN2 | SPTLC2 |  |  |
| DCTN5 | FBXO21 | SREK1 |  |  |
| DDI2 | FBXO31 | ST3GAL1 |  |  |
| DDX28 | FBXW2 | ST8SIA3 |  |  |
| DDX41 | FCHO2 | STC1 |  |  |
| DDX54 | FGF2 | STK40 |  |  |
| DDX6 | FJX1 | STOML1 |  |  |
| DGAT1 | FKBP14 | SUSD1 |  |  |
| DGCR2 | FKBP1A | TECPR2 |  |  |
| DGCR6L | FKBP4 | TEF |  |  |
| DGKA | FKBP9 | TENM4 |  |  |
| DHCR24 | FLCN | TMEM151B |  |  |
| DHX15 | FLNA | TMEM170A |  |  |
| DHX30 | FLNB | TMX4 |  |  |
| DHX38 | FMN1 | TRIB1 |  |  |
| DHX40 | FNDC3B | TRIM33 |  |  |
| DIMT1 | FNIP1 | TRIM35 |  |  |
| DIP2A | FOPNL | TSKU |  |  |
| DLG5 | FOXN2 | TXLNB |  |  |
| DMTN | FOXN3 | UBXN2A |  |  |
| DNAAF3 | FUT10 | VANGL2 |  |  |
| DNAJA3 | FUT11 | VAT1 |  |  |
| DNAJB2 | FXR1 | VPS37A |  |  |
| DNASE1L1 | FZD6 | WDR37 |  |  |
| DNMT1 | G2E3 | WDR45B |  |  |
| DNPEP | G3BP2 | WNT4 |  |  |
| DST | G6PD | WT1 |  |  |
| DSTN | GAA | XPNPEP3 |  |  |
| DVL2 | GALNT7 | XRRA1 |  |  |
| DYRK2 | GAN | ZDHHC22 |  |  |
| DZANK1 | GATA6 | ZDHHC5 |  |  |
| ECE1 | GATAD2A | ZNF391 |  |  |
| ECHDC2 | GATAD2B | ZNF431 |  |  |
| EDA2R | GCNT3 | ZNF488 |  |  |
| EED | GEMIN2 | ZNF550 |  |  |
| EEF1A1 | GFPT2 | ZNF638 |  |  |
| EEF1AKMT2 | GGCX | ZSCAN29 |  |  |
| EEF2 | GID4 |  |  |  |
| EFHC1 | GIT2 |  |  |  |
| EFNB3 | GLOD4 |  |  |  |
| EHMT1 | GLYR1 |  |  |  |
| EIF3A | GM2A |  |  |  |
| EIF4A1 | GNAI2 |  |  |  |
| EIF4B | GNAQ |  |  |  |
| EIF4G2 | GNB2 |  |  |  |
| EIF5B | GOLGA3 |  |  |  |
| EIF6 | GOLGA4 |  |  |  |
| ELAC2 | GOLGA8A |  |  |  |
| ELK4 | GOLGA8B |  |  |  |
| ELMSAN1 | GOLGA8IP |  |  |  |
| ELOA | GOLGA8J |  |  |  |
| ELOB | GPBP1L1 |  |  |  |
| ELOVL1 | GPR55 |  |  |  |
| ELP2 | GPX1 |  |  |  |
| EMC6 | GRAMD1B |  |  |  |
| EMC8 | GRAMD2B |  |  |  |
| EMD | GRAMD4 |  |  |  |
| ENKD1 | GRB2 |  |  |  |
| ENO1 | GRHPR |  |  |  |
| ENOPH1 | GSS |  |  |  |
| ENTPD4 | GSTM3 |  |  |  |
| ERVMER34-1 | GTF2A1 |  |  |  |
| EXOC5 | GTF2E1 |  |  |  |
| EXOC6B | GUF1 |  |  |  |
| EXOG | GULP1 |  |  |  |
| EXOSC10 | GXYLT1 |  |  |  |
| EXOSC2 | H3F3B |  |  |  |
| EZH2 | H3F3C |  |  |  |
| FAF1 | HAGH |  |  |  |
| FAM117A | HECTD1 |  |  |  |
| FAM129A | HIST1H2AM |  |  |  |
| FAM129B | HIST1H2BF |  |  |  |
| FAM160B2 | HIST2H2AC |  |  |  |
| FAM46A | HIST2H4B |  |  |  |
| FAR1 | HIVEP1 |  |  |  |
| FARSA | HMGA2 |  |  |  |
| FASN | HMGCR |  |  |  |
| FAU | HOXA13 |  |  |  |
| FBRS | HOXC8 |  |  |  |
| FBXL18 | HP1BP3 |  |  |  |
| FBXW2 | HPS6 |  |  |  |
| FEM1A | HSPA1B |  |  |  |
| FGB | IARS |  |  |  |
| FGFBP3 | IARS2 |  |  |  |
| FIS1 | IBTK |  |  |  |
| FIZ1 | ICAM1 |  |  |  |
| FKBP14 | IFIT3 |  |  |  |
| FKBP4 | IFITM1 |  |  |  |
| FLII | IFT22 |  |  |  |
| FLNA | IGSF8 |  |  |  |
| FLNB | IK |  |  |  |
| FLOT1 | IKZF2 |  |  |  |
| FURIN | IL6ST |  |  |  |
| FUT11 | ILF3 |  |  |  |
| FUZ | INCENP |  |  |  |
| FYCO1 | INSIG1 |  |  |  |
| G3BP1 | IPP |  |  |  |
| G3BP2 | IRGQ |  |  |  |
| GANAB | ITGA6 |  |  |  |
| GAPDH | ITGAV |  |  |  |
| GART | ITGB8 |  |  |  |
| GATAD2B | ITPR1 |  |  |  |
| GBP6 | JOSD1 |  |  |  |
| GCLM | KAT2B |  |  |  |
| GCN1 | KCNC4 |  |  |  |
| GEM | KDM2A |  |  |  |
| GEMIN4 | KDM3A |  |  |  |
| GFOD1 | KEAP1 |  |  |  |
| GHITM | KIAA0556 |  |  |  |
| GIGYF1 | KIAA1109 |  |  |  |
| GINS3 | KIAA1586 |  |  |  |
| GIT1 | KIAA1958 |  |  |  |
| GLCE | KIF1B |  |  |  |
| GLTP | KIF1BP |  |  |  |
| GLUL | KIF5B |  |  |  |
| GNB2 | KLHDC10 |  |  |  |
| GNL1 | KLHL14 |  |  |  |
| GNL2 | KLHL15 |  |  |  |
| GNS | KLHL18 |  |  |  |
| GOLGA3 | KLHL42 |  |  |  |
| GOLM1 | KMT2D |  |  |  |
| GOT2 | KMT5B |  |  |  |
| GPAM | LAMP2 |  |  |  |
| GPATCH4 | LARS |  |  |  |
| GPBP1 | LAX1 |  |  |  |
| GPR155 | LCOR |  |  |  |
| GPR26 | LDLR |  |  |  |
| GRB2 | LETM1 |  |  |  |
| GREB1L | LHFPL2 |  |  |  |
| GRIA4 | LILRA2 |  |  |  |
| GRIPAP1 | LONRF3 |  |  |  |
| GRPEL1 | LTBP3 |  |  |  |
| GTF2B | LUC7L3 |  |  |  |
| GTF2H3 | LYST |  |  |  |
| GTF2H4 | MAFK |  |  |  |
| GTF2I | MAN2A1 |  |  |  |
| GTF3C1 | MAP1B |  |  |  |
| GTF3C3 | MAP2K4 |  |  |  |
| GTPBP1 | MAP3K2 |  |  |  |
| GTSE1 | MAPK1 |  |  |  |
| H1F0 | MAST3 |  |  |  |
| H2AFV | MBD2 |  |  |  |
| H2AFX | MBNL1 |  |  |  |
| H2AFY | MCF2L2 |  |  |  |
| HADHA | MCL1 |  |  |  |
| HARS2 | MCOLN2 |  |  |  |
| HCFC1 | MDK |  |  |  |
| HCLS1 | MDM2 |  |  |  |
| HDHD5 | MED19 |  |  |  |
| HDLBP | MED29 |  |  |  |
| HEATR5A | MED7 |  |  |  |
| HECTD3 | MEF2D |  |  |  |
| HIC2 | METRN |  |  |  |
| HIPK1 | MEX3B |  |  |  |
| HIST1H3H | MFF |  |  |  |
| HIST1H4D | MFN1 |  |  |  |
| HIST2H4B | MIA3 |  |  |  |
| HIVEP2 | MKNK2 |  |  |  |
| HK2 | MMS19 |  |  |  |
| HLA-C | MOAP1 |  |  |  |
| HMGN1 | MORC3 |  |  |  |
| HMGXB3 | MPHOSPH10 |  |  |  |
| HNRNPA0 | MPP1 |  |  |  |
| HNRNPA1 | MRO |  |  |  |
| HNRNPA3 | MRPL19 |  |  |  |
| HNRNPC | MRPL39 |  |  |  |
| HNRNPH2 | MRPL53 |  |  |  |
| HNRNPK | MRPS16 |  |  |  |
| HNRNPM | MRPS21 |  |  |  |
| HOMER1 | MRPS23 |  |  |  |
| HOXA11 | MRS2 |  |  |  |
| HOXA5 | MSH3 |  |  |  |
| HOXC11 | MTF1 |  |  |  |
| HSD17B4 | MTMR1 |  |  |  |
| HSP90AA1 | MTMR10 |  |  |  |
| HSP90AB1 | MUC21 |  |  |  |
| HSP90B1 | MYC |  |  |  |
| HSPA1B | MYH10 |  |  |  |
| HSPBP1 | MYLIP |  |  |  |
| HSPE1-MOB4 | MYO1D |  |  |  |
| HUWE1 | MYO5A |  |  |  |
| HYOU1 | MYZAP |  |  |  |
| IFNAR2 | NABP2 |  |  |  |
| IGF1R | NACC2 |  |  |  |
| IGF2BP1 | NARF |  |  |  |
| IL2 | NCAPD2 |  |  |  |
| ILF2 | NCAPG2 |  |  |  |
| ILF3 | NCL |  |  |  |
| IMPA2 | NECAP1 |  |  |  |
| INO80B | NEMP1 |  |  |  |
| INTS5 | NF2 |  |  |  |
| IPO5 | NFATC2IP |  |  |  |
| IRS2 | NFYB |  |  |  |
| IRS4 | NIPA2 |  |  |  |
| ISM2 | NKAP |  |  |  |
| ITGA5 | NLK |  |  |  |
| ITGA9 | NLRP9 |  |  |  |
| IVD | NOL4L |  |  |  |
| JADE3 | NONO |  |  |  |
| JAG2 | NPM1 |  |  |  |
| JMJD6 | NPTN |  |  |  |
| KANSL3 | NPY4R |  |  |  |
| KARS | NRAS |  |  |  |
| KCNJ14 | NSF |  |  |  |
| KDM5C | NUCKS1 |  |  |  |
| KIAA0895 | NUFIP2 |  |  |  |
| KIAA1109 | NUGGC |  |  |  |
| KIF1C | NUP43 |  |  |  |
| KIF21B | OGFR |  |  |  |
| KLC2 | OPA3 |  |  |  |
| KLF2 | OR2A4 |  |  |  |
| KLHL26 | OR6V1 |  |  |  |
| KMT2A | ORAI2 |  |  |  |
| KMT2C | ORAOV1 |  |  |  |
| KNTC1 | ORC1 |  |  |  |
| KPNB1 | OSMR |  |  |  |
| KREMEN1 | OTUD3 |  |  |  |
| L3MBTL2 | OTUD7B |  |  |  |
| LAMB2 | PA2G4 |  |  |  |
| LAMP1 | PABPN1 |  |  |  |
| LARP1 | PAFAH1B1 |  |  |  |
| LARS | PAIP1 |  |  |  |
| LBR | PAPD5 |  |  |  |
| LDLR | PAPD7 |  |  |  |
| LETM1 | PARD6B |  |  |  |
| LIN28B | PARK7 |  |  |  |
| LIN7C | PARVG |  |  |  |
| LMNB1 | PAWR |  |  |  |
| LMOD3 | PAX9 |  |  |  |
| LONRF2 | PBLD |  |  |  |
| LRCH4 | PCBD2 |  |  |  |
| LRRC20 | PCMTD1 |  |  |  |
| LRRC58 | PDIK1L |  |  |  |
| LSM12 | PDZD8 |  |  |  |
| LUC7L3 | PEAK1 |  |  |  |
| LYRM7 | PELP1 |  |  |  |
| M6PR | PER2 |  |  |  |
| MAF1 | PFKFB4 |  |  |  |
| MAN2A2 | PFN1 |  |  |  |
| MAP2K7 | PGAM1 |  |  |  |
| MAP3K10 | PGAM4 |  |  |  |
| MAP4 | PGPEP1 |  |  |  |
| MAP7D1 | PHF19 |  |  |  |
| MARS | PHF23 |  |  |  |
| MAST3 | PHLPP2 |  |  |  |
| MAT2A | PHRF1 |  |  |  |
| MB21D1 | PHTF2 |  |  |  |
| MB21D2 | PIGO |  |  |  |
| MBOAT7 | PIK3AP1 |  |  |  |
| MCM3AP | PIK3CD |  |  |  |
| MCM5 | PIN4 |  |  |  |
| MCM7 | PIP5K1C |  |  |  |
| MCUR1 | PITPNA |  |  |  |
| MDH2 | PKNOX1 |  |  |  |
| MDM2 | PLA2G12A |  |  |  |
| MDN1 | PLEKHA1 |  |  |  |
| MED12L | PLXNA3 |  |  |  |
| MED15 | PLXND1 |  |  |  |
| MEGF8 | PMEPA1 |  |  |  |
| MEIS1 | PNO1 |  |  |  |
| METTL2A | POFUT1 |  |  |  |
| METTL2B | POLQ |  |  |  |
| MFN2 | POM121 |  |  |  |
| MFSD12 | PPCS |  |  |  |
| MGAT4B | PPIC |  |  |  |
| MGAT5 | PPIL1 |  |  |  |
| MIEF1 | PPP1CB |  |  |  |
| MINDY4 | PPP1R12C |  |  |  |
| MINOS1 | PPP1R37 |  |  |  |
| MKI67 | PPP1R3D |  |  |  |
| MKRN2 | PRKCSH |  |  |  |
| MLH3 | PRMT5 |  |  |  |
| MLLT1 | PRPS1 |  |  |  |
| MLX | PRR5 |  |  |  |
| MOB4 | PRRC2A |  |  |  |
| MPP2 | PRRC2B |  |  |  |
| MRM3 | PRRG4 |  |  |  |
| MRPL3 | PSMA7 |  |  |  |
| MRPL46 | PSMB3 |  |  |  |
| MRPS18B | PSMD3 |  |  |  |
| MRPS24 | PSMD5 |  |  |  |
| MSH6 | PSME3 |  |  |  |
| MSI2 | PTAR1 |  |  |  |
| MSL3 | PTBP1 |  |  |  |
| MSTO1 | PTEN |  |  |  |
| MTCH2 | PTGER4 |  |  |  |
| MTG1 | PTGES2 |  |  |  |
| MTHFD1L | PTPRJ |  |  |  |
| MTIF2 | PURG |  |  |  |
| MTR | PWP1 |  |  |  |
| MYH9 | PYCR2 |  |  |  |
| MYO10 | QSER1 |  |  |  |
| MYO5A | RAB23 |  |  |  |
| MYO9B | RAB3D |  |  |  |
| MYSM1 | RAB7B |  |  |  |
| MZT2B | RAB8B |  |  |  |
| N4BP2L2 | RAD21 |  |  |  |
| NABP2 | RAD51 |  |  |  |
| NADK2 | RAN |  |  |  |
| NASP | RANBP6 |  |  |  |
| NCAPD2 | RBBP5 |  |  |  |
| NCAPH | RBFOX2 |  |  |  |
| NCOA4 | RBL2 |  |  |  |
| NCOR1 | RBM10 |  |  |  |
| NCS1 | RBM14 |  |  |  |
| ND6 | RBM27 |  |  |  |
| NDST1 | RBM28 |  |  |  |
| NDUFA13 | RBM8A |  |  |  |
| NDUFV1 | RBMS2 |  |  |  |
| NDUFV3 | RBPJ |  |  |  |
| NEK1 | RECK |  |  |  |
| NEO1 | REL |  |  |  |
| NFE2L1 | RERE |  |  |  |
| NHLRC2 | REV3L |  |  |  |
| NHSL1 | REXO1 |  |  |  |
| NLE1 | REXO2 |  |  |  |
| NMD3 | RLIM |  |  |  |
| NME4 | RNF103 |  |  |  |
| NME6 | RNF168 |  |  |  |
| NME8 | RNF181 |  |  |  |
| NOA1 | RNF4 |  |  |  |
| NOC2L | RNF44 |  |  |  |
| NOL7 | RP2 |  |  |  |
| NOLC1 | RPAP1 |  |  |  |
| NOM1 | RPL23 |  |  |  |
| NOMO3 | RPL24 |  |  |  |
| NONO | RPL3 |  |  |  |
| NOTCH3 | RPL8 |  |  |  |
| NPAS2 | RPL9 |  |  |  |
| NPEPPS | RPLP0 |  |  |  |
| NRIP2 | RPRD2 |  |  |  |
| NSFL1C | RPS14 |  |  |  |
| NT5DC2 | RPS28 |  |  |  |
| NUBPL | RPS3A |  |  |  |
| NUCB1 | RPS4X |  |  |  |
| NUCKS1 | RRN3 |  |  |  |
| NUDC | RRP1 |  |  |  |
| NUDT3 | RSBN1 |  |  |  |
| NUFIP2 | RTKN2 |  |  |  |
| NUP54 | SASH1 |  |  |  |
| NUS1 | SCAF1 |  |  |  |
| NYNRIN | SCAF11 |  |  |  |
| OCRL | SELENOT |  |  |  |
| OPA3 | SERTAD2 |  |  |  |
| ORAI2 | SERTAD3 |  |  |  |
| ORC6 | SESN3 |  |  |  |
| P4HB | SETD5 |  |  |  |
| PA2G4 | SFPQ |  |  |  |
| PAGR1 | SFXN1 |  |  |  |
| PAN2 | SGK3 |  |  |  |
| PARP1 | SGPP1 |  |  |  |
| PARVB | SH2B3 |  |  |  |
| PBOV1 | SH3PXD2A |  |  |  |
| PCNT | SHE |  |  |  |
| PCYT1A | SIK1 |  |  |  |
| PDE12 | SIN3A |  |  |  |
| PDLIM2 | SKI |  |  |  |
| PDXK | SLC10A7 |  |  |  |
| PEG10 | SLC12A5 |  |  |  |
| PER1 | SLC15A1 |  |  |  |
| PFKL | SLC25A16 |  |  |  |
| PGAM1 | SLC25A32 |  |  |  |
| PGD | SLC25A36 |  |  |  |
| PHACTR4 | SLC25A4 |  |  |  |
| PHB | SLC25A5 |  |  |  |
| PHF23 | SLC33A1 |  |  |  |
| PHKA1 | SLC37A3 |  |  |  |
| PHKB | SLC39A14 |  |  |  |
| PHLPP1 | SLC39A6 |  |  |  |
| PHTF1 | SLC7A1 |  |  |  |
| PI4KA | SLC7A11 |  |  |  |
| PIN1 | SLC9A1 |  |  |  |
| PITHD1 | SLCO4A1 |  |  |  |
| PKD1 | SLX4 |  |  |  |
| PKM | SMAD3 |  |  |  |
| PLAGL2 | SMAD6 |  |  |  |
| PLD3 | SMAD7 |  |  |  |
| PLD6 | SMAP2 |  |  |  |
| PLOD1 | SMARCA5 |  |  |  |
| PLPBP | SMARCC2 |  |  |  |
| PLXNA1 | SMG1 |  |  |  |
| PMEPA1 | SMU1 |  |  |  |
| PMM2 | SNN |  |  |  |
| PNPT1 | SNRNP70 |  |  |  |
| POLDIP3 | SNRPD1 |  |  |  |
| POLG | SNX10 |  |  |  |
| POLR2A | SOGA1 |  |  |  |
| POLR3H | SOX11 |  |  |  |
| POLR3K | SOX4 |  |  |  |
| POM121 | SPATS2L |  |  |  |
| PPARGC1B | SPC24 |  |  |  |
| PPM1A | SPCS3 |  |  |  |
| PPM1G | SPOCK2 |  |  |  |
| PPP1R26 | SPRYD4 |  |  |  |
| PPP2R5A | SRFBP1 |  |  |  |
| PRADC1 | SRP68 |  |  |  |
| PRDX1 | SRPRA |  |  |  |
| PRDX4 | SSFA2 |  |  |  |
| PREPL | SSRP1 |  |  |  |
| PRKCD | STAT2 |  |  |  |
| PRMT7 | STRBP |  |  |  |
| PRPF31 | SUPT5H |  |  |  |
| PRPF8 | SUPT7L |  |  |  |
| PRPS1 | SURF6 |  |  |  |
| PRPS1L1 | SYMPK |  |  |  |
| PRR14L | SYNGAP1 |  |  |  |
| PRRC2A | SYNJ1 |  |  |  |
| PRRC2B | SZRD1 |  |  |  |
| PSMC2 | TACC1 |  |  |  |
| PSMC3IP | TAF12 |  |  |  |
| PSMC5 | TAF8 |  |  |  |
| PSMD8 | TANK |  |  |  |
| PSME3 | TATDN3 |  |  |  |
| PSMG1 | TBC1D8 |  |  |  |
| PTBP3 | TCOF1 |  |  |  |
| PTGR2 | TCP11L1 |  |  |  |
| PTRH2 | TECPR2 |  |  |  |
| PUF60 | TEF |  |  |  |
| PURA | TESK1 |  |  |  |
| PYGB | THAP10 |  |  |  |
| PYGO2 | TIRAP |  |  |  |
| R3HDM4 | TLN1 |  |  |  |
| RAB11FIP3 | TLR3 |  |  |  |
| RAB36 | TMEM115 |  |  |  |
| RAB3IP | TMEM184B |  |  |  |
| RAB40C | TMEM239 |  |  |  |
| RABL6 | TMEM33 |  |  |  |
| RAC3 | TMEM41A |  |  |  |
| RACGAP1 | TMEM44 |  |  |  |
| RAD51B | TMF1 |  |  |  |
| RASGRP3 | TNFRSF13C |  |  |  |
| RBM39 | TNPO1 |  |  |  |
| RBM42 | TNPO2 |  |  |  |
| RBMS2 | TOB1 |  |  |  |
| RC3H2 | TOB2 |  |  |  |
| RCC2 | TOR1B |  |  |  |
| RCE1 | TOR4A |  |  |  |
| RER1 | TPPP |  |  |  |
| REXO1 | TPT1 |  |  |  |
| REXO4 | TRAFD1 |  |  |  |
| RFTN1 | TRAM2 |  |  |  |
| RFWD3 | TRIM36 |  |  |  |
| RFX5 | TRIO |  |  |  |
| RGP1 | TRMT2B |  |  |  |
| RING1 | TSC1 |  |  |  |
| RIOK2 | TSPAN31 |  |  |  |
| RNASEH1 | TSPYL1 |  |  |  |
| RNF10 | TUBB6 |  |  |  |
| RNF11 | TULP4 |  |  |  |
| RNF111 | TWF1 |  |  |  |
| RNF115 | TXLNA |  |  |  |
| RNF14 | TXNDC15 |  |  |  |
| RNF167 | U2AF2 |  |  |  |
| RNF19B | UBE2Q2 |  |  |  |
| RNF216 | UBE2Z |  |  |  |
| RNF5 | UBXN4 |  |  |  |
| RNMT | UCK2 |  |  |  |
| RPL21 | UGDH |  |  |  |
| RPL27A | UHRF1BP1 |  |  |  |
| RPL28 | UNC5B |  |  |  |
| RPL36 | UQCRFS1 |  |  |  |
| RPL5 | UROD |  |  |  |
| RPLP0 | USF1 |  |  |  |
| RPLP2 | USP28 |  |  |  |
| RPN2 | UVRAG |  |  |  |
| RPS10 | VDAC2 |  |  |  |
| RPS14 | VHLL |  |  |  |
| RPS15 | VMA21 |  |  |  |
| RPS18 | VPS4B |  |  |  |
| RPS23 | VPS54 |  |  |  |
| RPS24 | WARS |  |  |  |
| RPS27L | WASL |  |  |  |
| RPS29 | WDR81 |  |  |  |
| RPS9 | WDR82 |  |  |  |
| RRAGD | WRNIP1 |  |  |  |
| RRM2 | XKR7 |  |  |  |
| RRP1B | XPO5 |  |  |  |
| RRS1 | XPO7 |  |  |  |
| RTN3 | XPR1 |  |  |  |
| RUNX2 | XRCC1 |  |  |  |
| RXRB | XRN1 |  |  |  |
| SALL2 | YIPF4 |  |  |  |
| SAP18 | YWHAE |  |  |  |
| SAV1 | ZADH2 |  |  |  |
| SCAP | ZBTB34 |  |  |  |
| SCD | ZBTB48 |  |  |  |
| SCIMP | ZBTB8B |  |  |  |
| SCYL1 | ZC3HAV1L |  |  |  |
| SDF4 | ZDHHC21 |  |  |  |
| SEC11A | ZDHHC24 |  |  |  |
| SEC16A | ZDHHC5 |  |  |  |
| SEMA4D | ZFC3H1 |  |  |  |
| SEPHS1 | ZFP62 |  |  |  |
| SEPHS2 | ZFYVE21 |  |  |  |
| SERPINH1 | ZIC5 |  |  |  |
| SESN2 | ZNF134 |  |  |  |
| SET | ZNF157 |  |  |  |
| SETD1B | ZNF17 |  |  |  |
| SETD6 | ZNF224 |  |  |  |
| SETD7 | ZNF24 |  |  |  |
| SF3B2 | ZNF264 |  |  |  |
| SF3B3 | ZNF267 |  |  |  |
| SGK3 | ZNF277 |  |  |  |
| SH2B1 | ZNF282 |  |  |  |
| SIPA1L2 | ZNF317 |  |  |  |
| SKAP2 | ZNF354B |  |  |  |
| SKI | ZNF383 |  |  |  |
| SLC10A7 | ZNF417 |  |  |  |
| SLC11A2 | ZNF430 |  |  |  |
| SLC16A1 | ZNF460 |  |  |  |
| SLC19A1 | ZNF492 |  |  |  |
| SLC1A4 | ZNF598 |  |  |  |
| SLC2A1 | ZNF607 |  |  |  |
| SLC35F5 | ZNF695 |  |  |  |
| SLC38A1 | ZNF721 |  |  |  |
| SLC39A13 | ZNF75A |  |  |  |
| SLC43A1 | ZNF772 |  |  |  |
| SLC4A2 | ZNF850 |  |  |  |
| SLC5A2 | ZNF98 |  |  |  |
| SLC6A8 | ZNRF3 |  |  |  |
| SLC7A5 | ZSCAN12 |  |  |  |
| SLK | ZSCAN18 |  |  |  |
| SMAD2 |  |  |  |  |
| SMC4 |  |  |  |  |
| SNRPD2 |  |  |  |  |
| SNRPD3 |  |  |  |  |
| SNX11 |  |  |  |  |
| SORD |  |  |  |  |
| SP3 |  |  |  |  |
| SPAG9 |  |  |  |  |
| SPECC1L |  |  |  |  |
| SPHK1 |  |  |  |  |
| SPIRE1 |  |  |  |  |
| SQSTM1 |  |  |  |  |
| SRCAP |  |  |  |  |
| SREBF1 |  |  |  |  |
| SRF |  |  |  |  |
| SRPRA |  |  |  |  |
| SRSF10 |  |  |  |  |
| SRSF9 |  |  |  |  |
| SS18L1 |  |  |  |  |
| ST3GAL2 |  |  |  |  |
| ST6GALNAC1 |  |  |  |  |
| STIP1 |  |  |  |  |
| STK10 |  |  |  |  |
| STK40 |  |  |  |  |
| STMN1 |  |  |  |  |
| STRN |  |  |  |  |
| STT3A |  |  |  |  |
| SUCO |  |  |  |  |
| SUGP2 |  |  |  |  |
| SUPT16H |  |  |  |  |
| SUPT5H |  |  |  |  |
| SURF2 |  |  |  |  |
| SYNE2 |  |  |  |  |
| SYNGR2 |  |  |  |  |
| SYNJ2BP |  |  |  |  |
| TAOK2 |  |  |  |  |
| TARBP2 |  |  |  |  |
| TARS2 |  |  |  |  |
| TBC1D15 |  |  |  |  |
| TBC1D22A |  |  |  |  |
| TBC1D2B |  |  |  |  |
| TBRG1 |  |  |  |  |
| TBX1 |  |  |  |  |
| TCF19 |  |  |  |  |
| TCHP |  |  |  |  |
| TEP1 |  |  |  |  |
| TET2 |  |  |  |  |
| TEX261 |  |  |  |  |
| TFAM |  |  |  |  |
| THAP6 |  |  |  |  |
| THOP1 |  |  |  |  |
| TIMELESS |  |  |  |  |
| TIMM10 |  |  |  |  |
| TIMM23 |  |  |  |  |
| TIMM50 |  |  |  |  |
| TIPIN |  |  |  |  |
| TIRAP |  |  |  |  |
| TKT |  |  |  |  |
| TLN1 |  |  |  |  |
| TM9SF3 |  |  |  |  |
| TMBIM6 |  |  |  |  |
| TMED9 |  |  |  |  |
| TMEM120B |  |  |  |  |
| TMEM131 |  |  |  |  |
| TMEM161A |  |  |  |  |
| TMEM170B |  |  |  |  |
| TMEM184B |  |  |  |  |
| TMEM2 |  |  |  |  |
| TMEM243 |  |  |  |  |
| TMEM50A |  |  |  |  |
| TMEM98 |  |  |  |  |
| TMEM9B |  |  |  |  |
| TMF1 |  |  |  |  |
| TMOD2 |  |  |  |  |
| TNFAIP1 |  |  |  |  |
| TNFAIP8L1 |  |  |  |  |
| TNFRSF12A |  |  |  |  |
| TNK2 |  |  |  |  |
| TNKS1BP1 |  |  |  |  |
| TNRC6A |  |  |  |  |
| TNRC6B |  |  |  |  |
| TOMM40 |  |  |  |  |
| TOMM40L |  |  |  |  |
| TOP1 |  |  |  |  |
| TOPBP1 |  |  |  |  |
| TOR2A |  |  |  |  |
| TPST2 |  |  |  |  |
| TRAF7 |  |  |  |  |
| TRAT1 |  |  |  |  |
| TRIM28 |  |  |  |  |
| TRIM44 |  |  |  |  |
| TRIM65 |  |  |  |  |
| TRIP6 |  |  |  |  |
| TROAP |  |  |  |  |
| TSC22D3 |  |  |  |  |
| TSEN54 |  |  |  |  |
| TSPAN3 |  |  |  |  |
| TSPAN33 |  |  |  |  |
| TSPAN6 |  |  |  |  |
| TTC36 |  |  |  |  |
| TTLL12 |  |  |  |  |
| TTYH3 |  |  |  |  |
| TUBA1B |  |  |  |  |
| TUBB |  |  |  |  |
| TUBB2A |  |  |  |  |
| TUBGCP4 |  |  |  |  |
| TULP4 |  |  |  |  |
| TUT1 |  |  |  |  |
| TWF2 |  |  |  |  |
| TXLNA |  |  |  |  |
| TYMS |  |  |  |  |
| UBA2 |  |  |  |  |
| UBA52 |  |  |  |  |
| UBAC2 |  |  |  |  |
| UBAP2L |  |  |  |  |
| UBE2S |  |  |  |  |
| UBE2Z |  |  |  |  |
| UBL7 |  |  |  |  |
| UBQLN1 |  |  |  |  |
| UBQLN2 |  |  |  |  |
| UBQLN4 |  |  |  |  |
| UBR5 |  |  |  |  |
| UBTF |  |  |  |  |
| UCP2 |  |  |  |  |
| UGGT2 |  |  |  |  |
| UPF3A |  |  |  |  |
| UQCRQ |  |  |  |  |
| URGCP |  |  |  |  |
| USP30 |  |  |  |  |
| USP40 |  |  |  |  |
| USP5 |  |  |  |  |
| UTP14C |  |  |  |  |
| UTRN |  |  |  |  |
| VAPB |  |  |  |  |
| VARS |  |  |  |  |
| VKORC1 |  |  |  |  |
| WAC |  |  |  |  |
| WASF2 |  |  |  |  |
| WDR26 |  |  |  |  |
| WDR4 |  |  |  |  |
| WDR5 |  |  |  |  |
| WDR91 |  |  |  |  |
| WEE1 |  |  |  |  |
| XPO7 |  |  |  |  |
| XRCC5 |  |  |  |  |
| XRCC6 |  |  |  |  |
| YBX1 |  |  |  |  |
| YRDC |  |  |  |  |
| YWHAE |  |  |  |  |
| YWHAZ |  |  |  |  |
| YY1 |  |  |  |  |
| ZBED4 |  |  |  |  |
| ZBTB4 |  |  |  |  |
| ZBTB45 |  |  |  |  |
| ZC3H18 |  |  |  |  |
| ZC3H7B |  |  |  |  |
| ZEB1 |  |  |  |  |
| ZFR |  |  |  |  |
| ZFX |  |  |  |  |
| ZFYVE21 |  |  |  |  |
| ZMAT5 |  |  |  |  |
| ZNF121 |  |  |  |  |
| ZNF202 |  |  |  |  |
| ZNF226 |  |  |  |  |
| ZNF264 |  |  |  |  |
| ZNF322 |  |  |  |  |
| ZNF347 |  |  |  |  |
| ZNF354A |  |  |  |  |
| ZNF362 |  |  |  |  |
| ZNF383 |  |  |  |  |
| ZNF48 |  |  |  |  |
| ZNF551 |  |  |  |  |
| ZNF557 |  |  |  |  |
| ZNF621 |  |  |  |  |
| ZNF664 |  |  |  |  |
| ZNF704 |  |  |  |  |
| ZNF792 |  |  |  |  |
| ZNF799 |  |  |  |  |
| ZNF8 |  |  |  |  |
| ZSCAN25 |  |  |  |  |
| ZSWIM8 |  |  |  |  |
| ZZEF1 |  |  |  |  |
