## Supplemental Table 3 for "Integrative transcriptomic analysis identifies miR-642a-5p as a regulator of *POFUT1* expression in colon cancer"

^*^ co‑last authors

Corresponding author:

Agnès Germot**^*1^**

**Supplemental Table 3** Target genes common at least to 2 selected miRNAs, extracted from MIENTURNET

| miRNA-target miRTarBase | | | | |
| --- | --- | --- | --- | --- |
| **hsa-miR-484** | **hsa-miR-92b-3p** | **hsa-miR-642a-5p** | **hsa-miR-3940-3p** | **hsa-miR-574-3p** |
| ADAM10 | ADAM10 | ATP2B4 | BCL2L1 | AMOTL1 |
| AMOTL1 | AP1B1 | BCL11B | BIRC5 | CLTC |
| AP1B1 | ARFGEF2 | CBX6 | C12orf49 | PEAK1 |
| ARFGEF2 | ARL6IP4 | CCDC171 | GPR75 | PIN1 |
| ARL6IP4 | ATF7IP | CLUAP1 | MLLT1 | RXRA |
| ATF7IP | ATP2A2 | DDI2 | ORAI2 |  |
| ATP2A2 | ATP2B4 | DHCR24 | RXRA |  |
| BAG6 | BAG6 | DKK3 | SCUBE3 |  |
| BBX | BBX | EEF2 | SH2B3 |  |
| BCAT2 | BCAT2 | ENTHD1 | STK10 |  |
| BCL2L1 | BCL11B | EXOC5 | TIMELESS |  |
| BIRC5 | CAPN15 | FBRS |  |  |
| C12orf49 | CAPRIN1 | GATAD2B |  |  |
| CAPN15 | CAPZB | GLUL |  |  |
| CAPRIN1 | CCDC171 | GPR75 |  |  |
| CAPZB | CD2AP | LARP1 |  |  |
| CBX6 | CLN8 | MDN1 |  |  |
| CD2AP | DDI2 | MRS2 |  |  |
| CLN8 | DKK3 | MTG1 |  |  |
| CLTC | EIF3A | MYO5A |  |  |
| CLUAP1 | ELOA | PAPD5 |  |  |
| DDI2 | ENTHD1 | PHF19 |  |  |
| DHCR24 | EXOC5 | PLXNA3 |  |  |
| EEF2 | EZH2 | **POFUT1** |  |  |
| EIF3A | FAM129A | SCUBE3 |  |  |
| ELOA | FAM46A | SMAD2 |  |  |
| EXOC5 | FAR1 | STK40 |  |  |
| EZH2 | FASN | TECPR2 |  |  |
| FAM129A | FBXW2 | TEF |  |  |
| FAM46A | FKBP14 | ZDHHC5 |  |  |
| FAR1 | FKBP4 |  |  |  |
| FASN | FLNA |  |  |  |
| FBRS | FLNB |  |  |  |
| FBXW2 | FUT11 |  |  |  |
| FKBP14 | G3BP2 |  |  |  |
| FKBP4 | GATAD2B |  |  |  |
| FLNA | GNB2 |  |  |  |
| FLNB | GOLGA3 |  |  |  |
| FUT11 | GRB2 |  |  |  |
| G3BP2 | HIST2H4B |  |  |  |
| GATAD2B | HSPA1B |  |  |  |
| GLUL | ILF3 |  |  |  |
| GNB2 | KIAA1109 |  |  |  |
| GOLGA3 | LARS |  |  |  |
| GRB2 | LDLR |  |  |  |
| HIST2H4B | LETM1 |  |  |  |
| HSPA1B | LUC7L3 |  |  |  |
| ILF3 | MAST3 |  |  |  |
| KIAA1109 | MDM2 |  |  |  |
| LARP1 | MRS2 |  |  |  |
| LARS | MYO5A |  |  |  |
| LDLR | NABP2 |  |  |  |
| LETM1 | NCAPD2 |  |  |  |
| LUC7L3 | NONO |  |  |  |
| MAST3 | NUCKS1 |  |  |  |
| MDM2 | NUFIP2 |  |  |  |
| MDN1 | OPA3 |  |  |  |
| MLLT1 | ORAI2 |  |  |  |
| MTG1 | PA2G4 |  |  |  |
| MYO5A | PAPD5 |  |  |  |
| NABP2 | PEAK1 |  |  |  |
| NCAPD2 | PGAM1 |  |  |  |
| NONO | PHF19 |  |  |  |
| NUCKS1 | PHF23 |  |  |  |
| NUFIP2 | PLXNA3 |  |  |  |
| OPA3 | PMEPA1 |  |  |  |
| ORAI2 | **POFUT1** |  |  |  |
| PA2G4 | POM121 |  |  |  |
| PGAM1 | PRPS1 |  |  |  |
| PHF23 | PRRC2A |  |  |  |
| PIN1 | PRRC2B |  |  |  |
| PMEPA1 | PSME3 |  |  |  |
| POM121 | RBMS2 |  |  |  |
| PRPS1 | REXO1 |  |  |  |
| PRRC2A | RPLP0 |  |  |  |
| PRRC2B | RPS14 |  |  |  |
| PSME3 | SGK3 |  |  |  |
| RBMS2 | SH2B3 |  |  |  |
| REXO1 | SKI |  |  |  |
| RPLP0 | SLC10A7 |  |  |  |
| RPS14 | SRPRA |  |  |  |
| SGK3 | SUPT5H |  |  |  |
| SKI | TECPR2 |  |  |  |
| SLC10A7 | TEF |  |  |  |
| SMAD2 | TIRAP |  |  |  |
| SRPRA | TLN1 |  |  |  |
| STK10 | TMEM184B |  |  |  |
| STK40 | TMF1 |  |  |  |
| SUPT5H | TULP4 |  |  |  |
| TIMELESS | TXLNA |  |  |  |
| TIRAP | UBE2Z |  |  |  |
| TLN1 | XPO7 |  |  |  |
| TMEM184B | YWHAE |  |  |  |
| TMF1 | ZDHHC5 |  |  |  |
| TULP4 | ZFYVE21 |  |  |  |
| TXLNA | ZNF264 |  |  |  |
| UBE2Z | ZNF383 |  |  |  |
| XPO7 |  |  |  |  |
| YWHAE |  |  |  |  |
| ZFYVE21 |  |  |  |  |
| ZNF264 |  |  |  |  |
| ZNF383 |  |  |  |  |
